# Molecular characterization of the N-terminal half of TasA during functional amyloid assembly and its contribution to *Bacillus subtilis* biofilm formation

**DOI:** 10.1101/2023.03.08.531758

**Authors:** Jesús Cámara-Almirón, Laura Domínguez-García, Nadia El Mammeri, Alons Lends, Birgit Habenstein, Antonio de Vicente, Antoine Loquet, Diego Romero

## Abstract

Biofilms are bacterial communities that result from a cell differentiation process that leads to the secretion of an extracellular matrix (ECM) by part of the bacterial population. In *Bacillus subtilis,* the main protein component of the ECM is the functional amyloid TasA, which forms a fiber-based scaffold that confers structure to the ECM. The N-terminal half of TasA is strongly conserved among *Bacillus* species and contains a protein domain, the amyloid core (AcTasA), which is critical for the formation of the amyloid architecture. In this study, we demonstrate that recombinantly purified AcTasA *in vitro* retains biochemical properties previously observed for the entire protein. Further analysis of the AcTasA amino acid sequence revealed two amyloidogenic stretches and a region of imperfect amino acid repeats, which are known to contribute to functional amyloid assembly. Biochemical characterization of these amyloidogenic stretches found in AcTasA revealed their amyloid capacity *in vitro*, contributing to the amyloid nature of AcTasA. Moreover, the study of the imperfect amino acid repeats revealed the critical role of residues D64, K68 and D69 in the structural function of TasA. *In vivo* and *in vitro* experiments with versions of TasA carrying the substitutions D64A, K68A, and D69A demonstrated a partial loss of function of the protein either in the assembly of the ECM or in the stability of the core and amyloid polymerization. Taken together, our findings allow us to better understand the polymerization process of TasA during biofilm formation and provide knowledge into the sequence determinants that promote the molecular behavior of functional amyloids.

## Introduction

Amyloids are proteins that have the ability to transition from monomers into insoluble fibers that share a common quaternary structure enriched in beta-sheets stacked in the so-called cross-beta architecture ^1^. These proteins have traditionally been studied in the context of human diseases, as several amyloid proteins and peptides are considered the causative agents of many protein misfolding disorders that lead to known neurodegenerative pathologies ^2, 3^. However, the beneficial role of amyloid proteins in many biological functions required for homeostasis is currently well established, which is why part of this broad family of proteins has been subclassified as functional amyloids ^3^. Alternative functions reported for these proteins in bacteria are antimicrobial, promotion of bacterial virulence in plants or providing a structural scaffold during biofilm formation (see ^4–6^ for extended reviews on the subject).

Biofilm formation is an intrinsic ability of bacterial species that allows their association in communities in which bacterial cells are closely linked to each other by the secretion of a self-produced extracellular matrix (ECM) ^7^. The ECM is mostly comprised of extracellular polysaccharides, proteins and other molecules that allow for a more efficient interaction of the microbial community with the environment and act as a shield that protects cells against external stressors ^7, 8^. Functional amyloids and other proteinaceous filaments play a key role in the assembly of these complex multicellular communities, providing mainly structural support. In species from the *Bacillus* genus, the main protein component of biofilms is TasA ^9^, which differs between the two phylogenetically distinct groups of *Bacillus*, the *subtilis* group and the *cereus* group ^10^. TasA, in its monomeric form, is a stable soluble globular protein that is able to transition during biofilm formation into filaments that exhibit typical amyloid features (resistance to protease degradation or enrichment in the beta-sheet secondary structure) ^10–12^. TasA fibers have been shown to form *in vivo*, by direct observation in biofilm samples ^12^, and *in vitro,* in protein extracted from biofilm samples in a pre-aggregated conformation, where the aggregation properties and morphology of the native filaments have been studied ^13–16^, or in heterologously expressed recombinant protein purified from *E. coli*, where the structural properties of the protein and fibers have been characterized ^10, 11^. The assembly of TasA is assisted by TapA, a two-domain partially disordered accessory protein ^17^ that is also essential for biofilm formation, which promotes TasA polymerization and anchors the fibers to the cell surface ^10, 18^. The main biological function of these fibers in biofilms is structural, creating a scaffold that provides integrity to the ECM ^12^. However, the protein plays other biologically relevant roles, contributing to membrane stability and dynamics or acting as a developmental signal that maintains a motile subpopulation within biofilms ^19, 20^.

The formation of the amyloid fold is not a trait exclusively restricted to amyloid proteins but is more a universal property of polypeptide chains when certain physicochemical conditions are achieved ^1^. In amyloids, this specific folding pattern is favored, which is partly due to the amino acid context where certain regions, the presence of different amino acid patterns (such as imperfect repeats, especially in bacterial functional amyloids ^21, 22^) or the appearance of specific amino acids in certain conformations determine the amyloid propensity of a given protein under physiological conditions ^23^. These regions have been studied extensively in amyloid proteins, as they form what is known as the amyloid core of the protein, the part of the protein that contributes more to the cross-beta structure and is more tightly folded ^24^, conferring the fibers their unique physicochemical properties.

In previous works ^10^, we studied the structural properties of TasA in its fibrillar form, where an amyloid core, comprising mostly the N-terminal half of the protein, was delimited. In the present work, we have revealed some of the sequence determinants of this amyloid core that promote the biochemical behavior of TasA. By combining bioinformatics, mutational analysis and structural, biochemical, and biophysical studies, we defined specific residues critical for the structural function of TasA in the ECM. All these results allow for a better comprehension of the structural function of TasA during ECM production and biofilm formation and thus improve our understanding of the role of functional amyloids and other filamentous proteins in the persistence and interaction of bacteria in and with the environment.

## Results

### TasA contains an N-terminal amyloid core that retains amyloid properties

Solid-state nuclear magnetic resonance spectroscopy analysis (SSNMR) on assembled recombinant TasA fibers allowed us to delimitate a rigid amyloid core (AcTasA) of 110 amino acids that extends from residues K35 to K144 ^10^ (Fig 1 residues labeled in red). AcTasA is located in the N-terminal half of the protein, which is the region of TasA that is highly conserved across different *Bacillus* species (Fig 1). Based on these findings, we hypothesized a relevant contribution of this region to the biochemical and biophysical properties of the protein and to the final architecture of the TasA fiber and functionality. We analyzed the properties of the AcTasA sequence using two complementary bioinformatic approaches. First, we searched for amino acid repeats within the sequence, which have been demonstrated to be involved in the polymerization process of other amyloid proteins, including bacterial functional amyloids ^21, 22, 25, 26^; second, we looked for aggregation-prone regions, also known as amyloidogenic stretches, in AcTasA. These tools revealed the presence of i) two amyloidogenic regions that extend from L78 to G90 (called here the segment LG-13) and from D104 to G117 (called DG-14) (Fig 1, labeled, respectively, in blue and green) and ii) a sequence of imperfect amino acid repeats in which the sequence KDxxFxxxxxxLxxKExxxxxNxxxxKxxxGxxxx is repeated twice and extends from K35 to S101 (Fig 1 underlined in cyan).

**Fig 1.**
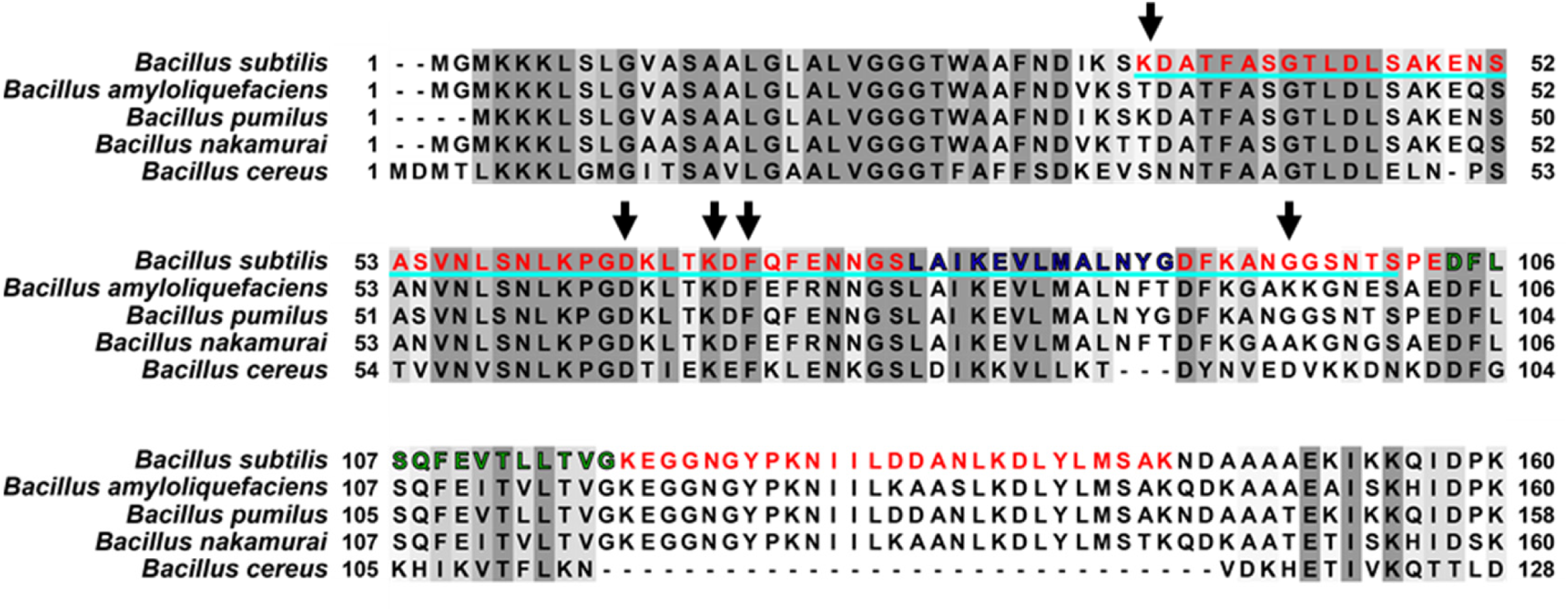
The N-terminal Domain of TasA Revealed Amyloidogenic Stretches and Imperfect Amino Acid Repeats. Sequence Comparison of the N-terminal Domain of TasA between Different Species of *Bacillus*. The different features detected by bioinformatic tools are labeled over the sequence. The gray colors indicate different degrees of sequence conservation. The region corresponding to the amyloid core of TasA is labeled in red (from K35 to K144). The two amyloidogenic stretches are labelled in blue (L78 to G90, LG-13) and green (D104, G117, DG-14). The sequence underlined in cyan corresponds to the imperfect amino acid repeat in which the sequence KDxxFxxxxxxLxxKExxxxxNxxxxKxxxGxxxx is repeated twice in the core. Arrows indicate residues that have been analyzed by site-directed mutagenesis.

To experimentally corroborate the predicted amyloid properties of this region, AcTasA, heterologously expressed in *E. coli*, was purified to homogeneity and maintained soluble in monomeric form in a 1% acetic solution ^10^ prior to studying the polymerization on buffer at physiological pH (20 mM Tris, 50 mM NaCl at pH 7). Transmission electron microscopy (TEM) analysis of AcTasA in buffer after 1 week of incubation revealed filaments with a bundle-like supramolecular organization that resembled those formed by the entire TasA protein (Fig 2A, top and bottom micrographs, respectively). The bundle widths varied from ∼5 to ∼25 nm with a tendency to stack themselves to form larger objects. The smallest molecular entity observable, presumably protofibrils, averaged ∼3.5 nm wide. Given the ability of AcTasA to aggregate and form filament-like structures, we further characterized the kinetics of the aggregation process over time using dynamic light scattering (DLS), a noninvasive technique that allows us to monitor the size distribution of the aggregates found in the sample based on the signal intensity associated with each hydrodynamic radius (R_H_). The absence of major changes in the size distribution of the population during the time studied and the mean R_H_ of ∼10 nm of the population immediately after buffer exchange (time 0 h), which is far above what is attributed to WT TasA ^10, 13^, were two complementary pieces of evidence suggesting that polymerization of AcTasA occurs rapidly upon buffer exchange to neutral pH (Fig 2B). Two populations of similar abundance values and mean R_H_ values of ∼12 nm and ∼22 nm were observed at 72 h, indicative of a transition into larger aggregates. According to this dynamic, a more homogeneous size distribution of aggregates with a mean R_H_ of ∼25 nm dominated at 144 h (Fig 2B, middle and bottom graphs). Overall, this set of experiments confirms that AcTasA rapidly transitions at pH 7 from monomers to aggregates that exhibit an unequivocal fibrillar form. Next, we wondered whether these AcTasA assemblies exhibited amyloid biochemical features similar to those of full-length WT TasA. Amyloid proteins exhibit unique tinctorial properties as they bind to specific dyes. One such dye is thioflavin-T (ThT), which upon binding to the beta-sheets naturally present in amyloid samples, undergoes a shift in the fluorescence emission maximum that can be used to estimate the gain of beta-sheet secondary structure; this occurs concomitantly with the transition from the monomeric to the fibrillar form ^27^. ThT binding assays of purified AcTasA showed a progressive increase in the fluorescence intensity of ThT at the amyloid-specific wavelength over time in a concentration-dependent manner (Fig 2C). Moreover, this ThT binding kinetics was shown in a polymerization curve similar to that observed for WT TasA *in vitro* and other amyloid proteins ^10, 28–30^.

**Fig 2.**
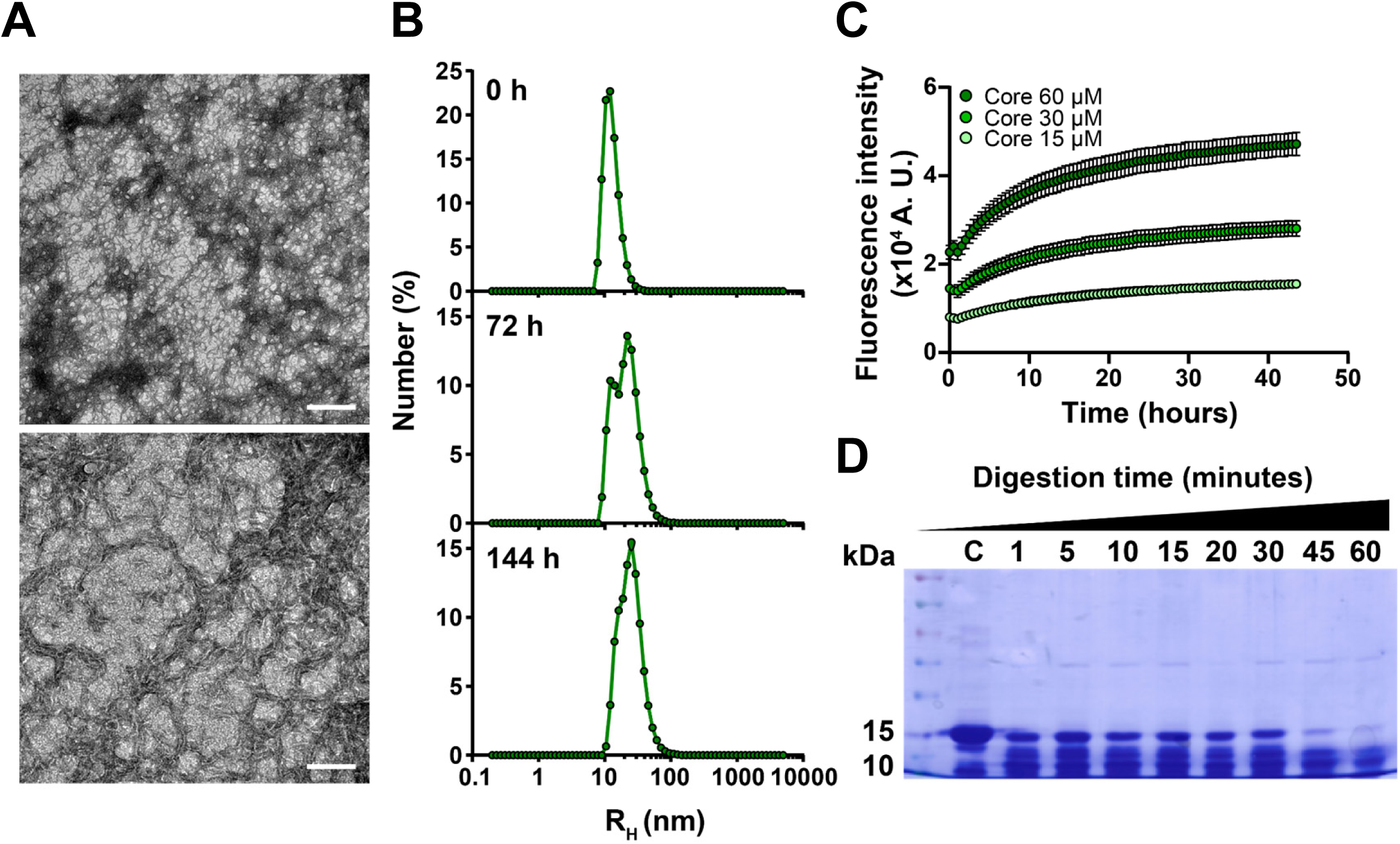
The TasA Amyloid Core Region Exhibits Amyloid Properties *In Vitro*. (A) Transmission electron micrographs of negatively stained AcTasA samples show dense bundles of fibrillar assemblies. The scale bars are 500 nm (top) and 100 nm (bottom). (B) Aggregation kinetics of AcTasA as measured by DLS at different time points. (C) Kinetics of fluorescence emission at 480 nm of AcTasA samples indicates ThT binding properties. Experiments were performed at different protein concentrations. Average values of three independent experiments are represented. Error bars indicate the SEM. (D) Coomassie stained SDS‒PAGE gel of the AcTasA samples digested with proteinase K at different time points. The first lane indicates the molecular marker. Each lane contains a sample corresponding to a specific digestion time.

As mentioned above, the amyloid core is the part of the protein most intensely folded in the final tridimensional structure of the fiber, and therefore, it exhibits remarkable physicochemical robustness, including resistance to proteases ^31^. In our previous work, AcTasA was detected after TasA fibrils were subjected to partial proteinase K digestion ^10^. Thus, we wondered if the aggregation properties of isolated AcTasA preserved the structural fold that confers TasA fibers their resistance against proteolytic activity. In a similar protease digestion experiment, polymerized AcTasA resisted treatment with proteinase K even 1 h after incubation, as shown by the ∼13 kDa band resolved in the comassie-stained SDS‒PAGE gel of the digestion samples compared to the untreated control (Fig 2D, last lane on the right and lane labeled with a “C”, respectively).

### Amyloidogenic features of the N-terminal half of TasA contribute to the molecular behavior of AcTasA

Our previous findings demonstrated that AcTasA exhibits biochemical features similar to those observed for the whole TasA protein and proved that these properties are retained in the N-terminal half. Thus, we asked if the features previously predicted in our *in silico* analysis (Fig 1) were relevant to defining the structure or functionality of TasA. Our bioinformatic predictions of amyloidogenic stretches consistently showed two regions of AcTasA, designated LG-13 and DG-14, with a high tendency for aggregation (Fig 3A) (see Fig 1, labeled in blue and green, respectively). To study whether these two regions contribute to biofilm formation and have any impact on the structural function of TasA, we performed a mutagenesis analysis in which either of these two stretches was deleted from the protein sequence. As described in a previous work ^19^, to avoid undesirable effects when manipulating the *tasA* gene in the endogenous operon, these experiments were performed using a strain lacking the whole *tapA-sipW-tasA* operon and complemented with a version of the operon containing the mutated versions of *tasA* and integrated in the neutral *lacA* locus. Complementation of the Δ*(tapA-sipW-tasA)* strain with an operon containing a native version of TasA rescued the wrinkled phenotype associated with the WT strain in solid and liquid cultures (Fig 3B, left images). However, complementation with the operon containing a TasA version lacking amino acids from E82 to N88 (Δ82-88, LG-13 region) or from Q108 to V116 (Δ108-116, DG-14 region) failed to restore the biofilm formation phenotype (Fig 3B, middle and right images). To check if these mutated versions of TasA were stable, we performed a biofilm fractionation assay followed by protein precipitation and Western blot analysis using anti-TasA antibodies. A strong anti-TasA reacting band was observed in samples from WT or the control strains, but the signal was absent in samples of the strains carrying the mutated alleles (Fig 3C). This result, along with the fact that the two strains bearing the mutated versions of TasA showed the same phenotype, suggested that the deletion of these regions renders the protein unstable, which most likely leads to the degradation of the protein by quality control proteases.

**Fig 3.**
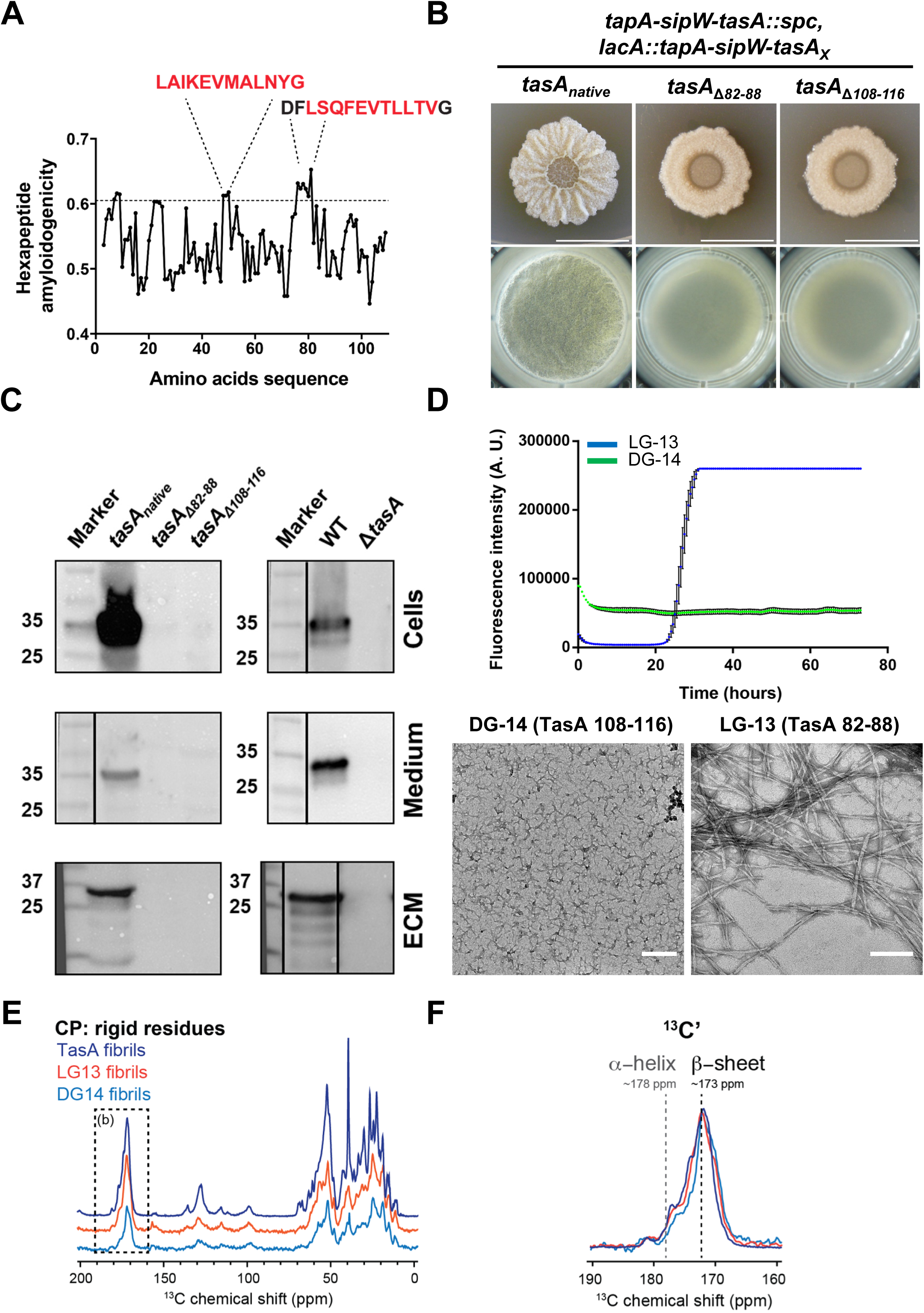
Amyloidogenic Regions are Important for Protein Folding and Contribute to the Amyloid Properties of AcTasA. (A) The hexapeptide amyloidogenicity profile of AcTasA as predicted by MetAmyl reveals two amyloidogenic stretches found in this sequence. (B) Mutagenesis analysis of AcTasA in which the predicted amyloid stretches were deleted from the TasA sequence and the mutated alleles were introduced in a Δ*(tapA-sipW-tasA)* background. The images show the phenotypes of *B. subtilis* biofilms grown in solid and liquid biofilm-inducing media of the strains carrying either the native allele (unmodified) or the alleles carrying the deletions. Scale bars = 1 cm. (C) Western blot using an α-TasA antibody of protein extracts corresponding to the different biofilm fractions (cells, medium and matrix) of the strains carrying the abovementioned alleles. Immunoblot images have been cropped and spliced for illustrative purposes. Black lines over the blot images delineate boundaries of immunoblot splicing. In all cases, all the slices shown were derived from a single blot. (D) Top. Kinetics of ThT fluorescence emission at 480 nm of the synthetic peptides corresponding to the amyloid stretch predicted. The average of three independent experiments is shown. Error bars indicate the SEM. Bottom. Transmission electron microscopy micrographs of negatively stained LG-13 and DG-14 synthetic peptides at the final experimental time-point. Scale bar = 200 nm. (E) ^1^D ^13^C cross-polarization spectra of TasA fibrils (dark blue) and the assemblies formed by peptides LG-13 (red) and DG14 (light blue). (F) Zoomed region of the 1D 13C cross-polarization spectra shown in E between 160 and 190 ppm.

Complementarily, we studied the amyloidogenic behavior of these two amino acids stretches *in vitro* using synthetic peptides. None of the peptides showed ThT binding activity (β−sheet formation) during the first 20 hours of incubation. However, after this latent phase, the LG-13 peptide exhibited an exponential increase in the fluorescence signal, reaching saturation a few hours later (Fig 3D, blue). Transmission electron microscopy analysis at the end of the ThT experiment revealed molecular entities of small size without any significant organization for the DG-14 peptide (Fig 3D, left image). Abundant fibrillar assemblies made of bundles comprised of several protofilaments (∼5 nm diameter each) ranging between 6 and 14 nm in diameter, which is a molecular organization very similar to the full-length mature TasA assemblies observed *in vitro*, characterized samples of the LG-13 peptide. We further studied the rigid core of the self-assembled fibrils made of LG-13 or the molecular entities of DG-14 TasA peptides by solid-state NMR spectroscopy. Using magic-angle spinning NMR, we measured 1D dipolar-based ^13^C cross-polarization (CP) spectra to assess the secondary structure conformation of TasA subunits along the fibril axis (Fig 3E). Both peptides showed strong and sharp ^13^C NMR signals (Fig 3E), indicating that the structural core of these fibrils is made of well-ordered and rigid subunits. We compared those 1D ^13^C spectra to that of full-length TasA fibrillated *in vitro* ^10^. Overall, the spectral envelopes of the three samples are quite similar, indicating a very similar secondary conformation. In particular, the backbone carbonyl ^13^C atoms accurately probe the conformation of amino acids using NMR chemical shifts. Indeed, β-sheet carbonyl chemical shifts are ∼173 ppm, while α-helical residues are found at ∼178 ppm (using a DSS-based NMR scale) (Fig 3F). Similar to full-length TasA, both LG-13 and DG-14 peptides form β-rich fibrils. Overall, SSNMR analysis suggests that LG-13 and DG-14 are amyloidogenic segments of TasA that maintain the same secondary structure conformation, and they also have the ability to self-assemble into fibrils formed by well-ordered subunits associated with low structural polymorphism.

### AcTasA contains an imperfect amino acid repeat with specific amino acids critical for protein functionality

Repeated regions within amyloid proteins have been demonstrated to contribute to the assembly of amyloid filaments ^21, 22, 32^, and an imperfect amino acid repeat, KDxxFxxxxxxLxxKExxxxxNxxxxKxxxGxxxx, which is repeated twice, was predicted within the AcTasA region (Fig 1, cyan line). To explore the contribution of this region to the structure and functionality of TasA, we performed a mutational analysis by alanine scanning on selected amino acids K35, D36, D64, K68, D69, F72 and G96, which showed different levels of conservation (Fig 1, arrows). The strain expressing the native *tasA* allele was unaffected in its ability to form a WT-like biofilm in solid or liquid medium (Fig 4A). Strains expressing some of the mutated alleles (K35A, D36A; F72A or G96A) also displayed the characteristic wrinkled morphology of *B. subtilis* biofilms (Fig 4A). However, the strains that expressed the D64A, K68A, and D69A alleles failed to fully restore the WT biofilm formation phenotype (Fig 4A).

**Fig 4.**
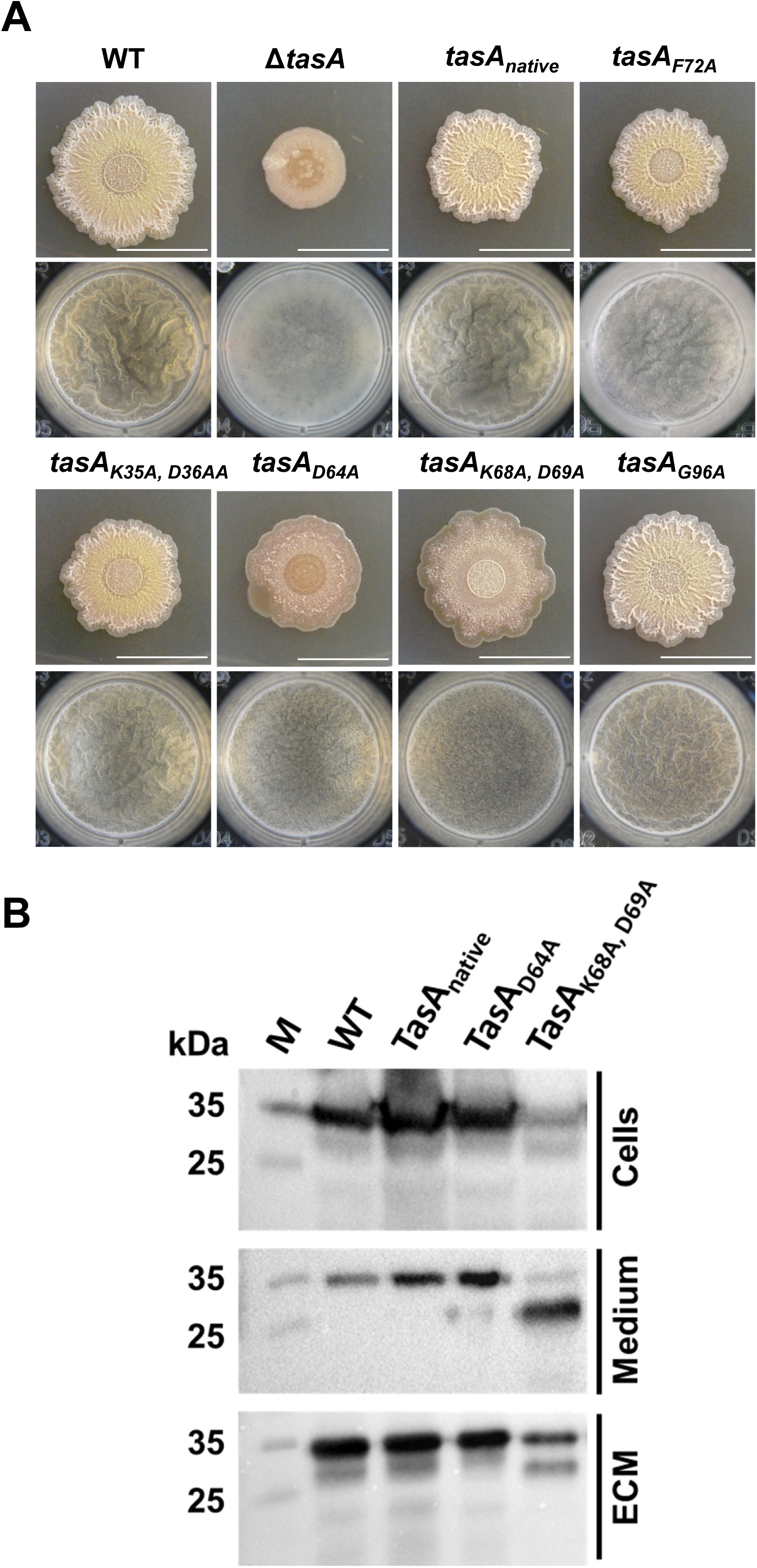
Mutagenesis Analysis of the Imperfect Amino Acid Repeat Reveals Amino Acids Essential for Biofilm Formation. (A) Alanine scanning site-directed mutagenesis analysis of AcTasA in which specific amino acids from the imperfect repeat region of AcTasA were substituted from the TasA sequence and the mutated alleles were introduced in a Δ*(tapA-sipW-tasA)* background. The images show the phenotypes of *B. subtilis* biofilms grown in solid and liquid biofilm-inducing media of the strains carrying either the native allele (unmodified) or the alleles carrying the substitutions. Scale bars = 1 cm. (B) Western blot using an α-TasA antibody of protein extracts corresponding to the different biofilm fractions (cells, medium and matrix) of the strains carrying the abovementioned alleles that were affected in colony morphology or pellicle formation. Immunoblot images have been cropped for illustrative purposes.

Apart from its structural role in fiber assembly and biofilm formation, the presence of TasA in the cell membrane plays an important role in bacterial physiology, maintaining cell membrane stability and preventing bacterial cell death ^19^. Thus, we wondered if the phenotypes that we observed in these two strains carrying the TasA variants could somehow be related to this physiological function of TasA in the cell membrane. Confocal microscopy analysis of strains expressing the *tasA* alleles that were fused to mCherry and staining with membrane dye showed a homogenous distribution of the TasA-related fluorescence signal across the cell membrane except in the poles, where it formed two small foci. The same distribution of TasA was observed in all samples, regardless of the mutations introduced (S1 Fig); thus, we discarded any effect in the observed phenotype that could be attributed to physiological alterations caused by a mislocalization of TasA.

We then speculated that this partial loss of phenotype could be related to the instability of the mutated TasA alleles due to the alanine substitutions. Biofilm fractionation assay from pellicles followed by immunoblot analysis using the anti-TasA antibody (Fig 4B) showed the presence of an anti-TasA reacting band in all the fractions from the WT strain, the strain carrying the native TasA allele or the strain carrying the TasA D64A allele. However, a weaker signal was observed in fractions of cells expressing the TasA K68A and D69A versions. In addition, two well-defined reacting bands appeared in the medium and ECM fractions of samples from this strain, which suggests interference of the amino acid substitution with protein processing. Nonetheless, we concluded from this experiment that the TasA version of the mutants with an altered biofilm formation phenotype is sufficiently stable to be detected in protein extracts; thus, the mutant phenotype should not be explained by the instability and degradation of TasA. TEM analysis of 48 h pellicles of these two mutated strains showed a network of branched fibers decorating the surfaces of cells expressing the native TasA or the TasA D64A alleles (S2 Fig, top and second from the top micrograph panels). In contrast, straight fibrillar structures were scarcely observed in cell preparations of the strain expressing the TasA K68, D69A allele (S2 Fig).

Thus, it appeared that the structural function of TasA in the formation of the fibers and required for biofilm formation was compromised in this strain. We performed an *in silico* analysis in which the mutated proteins were modeled and compared to the WT protein (Fig 5). The TasA WT structure had to be predicted and used as a reference when comparing the variant proteins, although the structure of the TasA monomer has already been experimentally determined and deposited in the Protein Data Bank ^11^. Differences between the modeled WT TasA predicted by AlphaFold and the crystallographic model were found in the loop between K118 and Y124, which was not modeled in the crystal structure, or in some regions that are modeled differently in the predicted WT structure, such as the first amino acids or the absence of beta-sheet β7 (S3 Fig). We represented the different protein models using the crystal structure as a reference, using the same schematic representation of the different structural features of the protein that was used in another study ^11^ (S3 Fig). When the models of the mutated versions of TasA and WT TasA were compared (Fig 5A and B), some noticeable structural differences were found: i) TasA D64A showed the disruption of the β6 beta-sheet by a random coil conformation adopted by residues from K126 to D130 and ii) the appearance of a new alpha-helix, labeled α5’, from residues G175 to T177 (Fig 5A). The disruption of the alpha-helix α5 was the only difference between the model of TasA K68A, D69A and the TasA WT model (Fig 5B). To further analyze these structural differences, we additionally modeled the possible hydrogen bonds within each protein structure. The D64 residue present in the loop between β2 and β3 in the WT protein forms five hydrogen bonds with neighboring amino acids: two with K61 and one with F200 (Fig 5C top, left image). The substitution of this residue seems to interfere with one of the hydrogen bonds formed with K61 (Fig 5C top). This single change seems to perturb the structure of the protein and might explain the structural differences when compared to the TasA WT model. In the case of TasA K68A and D69A, the K68 residue is bound to M196 from β8 through two hydrogen bonds (Fig 5C bottom). The substitution of these amino acids preserves the hydrogen bonds connecting the two beta-sheets by A68, but an additional bond is formed between A69 and G42. This change in the hydrogen bond configuration might result in the loss of α5, whose residues are modeled as random coils in the predicted structure of this TasA variant.

**Fig 5.**
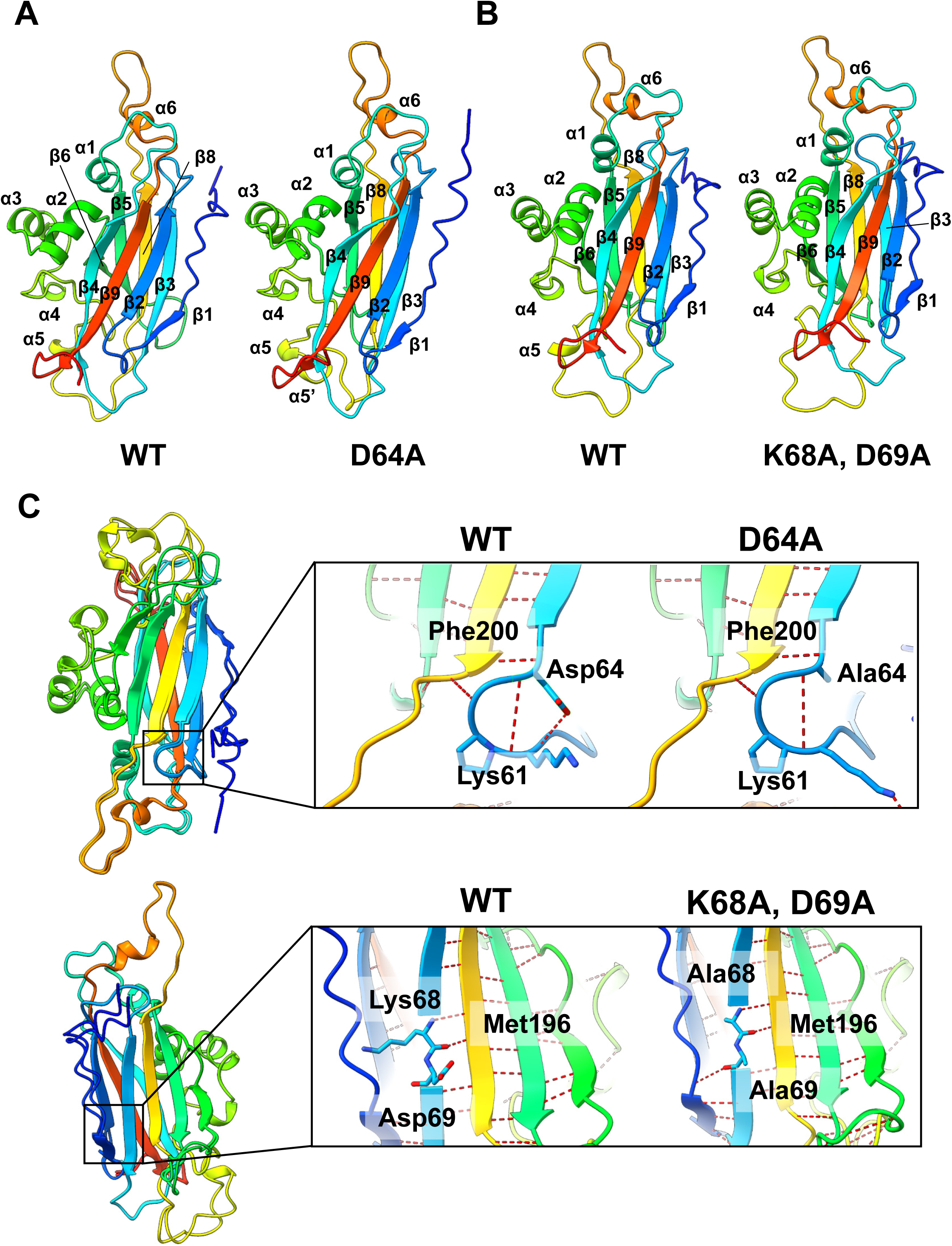
Structure Predictions and Comparison of the WT TasA and Variant Proteins. (A) Structure comparison between the TasA model and the D64A allele. The numbering in the scheme follows the representation described for the crystal structure of TasA in its monomeric form published by Diehl et al. ^11^. The coloring scheme indicates the position of the residue in the sequence of the protein, where blue indicates the N-terminus and red indicates the C-terminus. Warmer colors indicate proximity to the C-terminal end. (B) Structure comparison between the TasA model and the K68A and D69A alleles. (C) Superimposition of the WT – D64A models (top) and WT – K68A and D69A models (bottom) and a detailed view of the hydrogen bond analysis of the mutated proteins.

### Molecular characterization of the TasA variants revealed two different alterations in the core of the mutated proteins

Three residues of the AcTasA core, D64, K68 and D69, are relevant for the functionality of TasA, and the predicted structural changes of TasA associated with their substitution explain their high degree of conservation within the AcTasA sequence (see Fig 1). Thus, we reasoned that these amino acids participate in the polymerization of TasA. The two versions of TasA were heterologously expressed and purified, and their molecular properties were studied *in vitro* using the protocols previously optimized for TasA, TapA or AcTasA ^10^. The structural changes observed for these two versions of TasA were suggestive of alterations at the level of structural stability compared to the WT TasA. We used circular dichroism (CD) to measure the mean residue ellipticity (MRE) at 222 nm, a representative wavelength of the protein secondary structure, as a function of temperature (S4 Fig) to follow the unfolding of the proteins and investigate differences associated with the specific amino acid substitutions. Interestingly, the three proteins showed a high degree of thermostability. The thermal scan of WT TasA showed gain of secondary structure upon heating (S3 Fig), a thermal behavior that is distinct compared to that of globular proteins, especially at high temperatures, where heating normally induces protein unfolding and, therefore, an increase in the MRE. However, similar to what has been reported for other amyloid proteins ^33^, a decrease in the MRE as a function of temperature was recorded instead of reaching a completely unfolded state at high temperatures. TasA D64A, despite the structural alterations, exhibited a similar thermal behavior as that observed for TasA WT, showing enrichment in the secondary structure with increasing temperature and a higher signal at 222 nm, suggesting a lower proportion of secondary structure, especially in the range of 60 °C, compared to the WT protein. Major differences were evident between TasA K68A and D69A and TasA WT. A slight gain of secondary structure upon heating was observed in comparison to the other versions of TasA; however, the signal variations were very low compared to the other versions of TasA, indicating that the protein does not suffer profound changes in secondary structure at this wavelength with increasing temperature; therefore, changes in the thermal stability can be deduced.

To test whether these differences in structure and stability are translated into differences in the amyloid behavior of protein and fiber formation, we analyzed the aggregation kinetics of the two alleles using DLS. TasA D64A showed a mean R_H_ of ∼1.6 nm at the beginning of the experiment, which was slightly smaller than that reported for TasA WT ^10^. However, the most representative population in the TasA D64A sample reached a mean R_H_ between ∼7.8 and ∼9 nm after 72 h of incubation and finally reached a mean size between ∼9 and ∼10 nm after 144 h (Fig 6A left). The kinetics of aggregation of TasA K68A and D69A were, however, substantially different, given that the mean size distribution of the particles present in the sample was ∼2 nm at t = 0 h and remained nearly invariable over time (Fig 6A right). This finding suggested that the aggregation kinetics of this protein are slower than those observed for the WT protein or the TasA D64A allele, indicating a defect of TasA K68A and D69A in aggregation. We next measured the characteristic beta-sheet enrichment of amyloid proteins in these molecular assemblies by ThT binding experiments. The dynamics of fluorescence emission of TasA D64A (Fig 6B blue) at the amyloid-specific wavelength were similar to those of TasA WT (Fig 6B gray), indicating a similar beta-sheet content. However, the fluorescent signal of TasA K68A and D69A saturated earlier than that of the TasA WT or TasA K68A and D69A proteins, even at the higher protein concentration used in the experiment, and the maximum intensity of the signal was, overall, nearly three times lower than that observed for the TasA WT or TasA D64A proteins (Fig 6B red), suggesting limited beta-sheet enrichment. Morphological characterization by TEM of the molecular assemblies formed by the two mutated variants showed a tendency to form large aggregates compared to TasA WT (Fig 6C). The aggregates of TasA D64A were composed of associated bundles of filaments of different sizes, from ∼5 nm to hundreds of nanometers. Some individual fibrillar entities could also be observed within these bundles, with an average size between ∼1.5 and ∼2.5 nm wide and with variable length (Fig 6C center panel). TasA K68A and D69A, however, formed large and densely packed aggregates with individual fibrillar entities that were hardly visible (Fig 6C right panel). The partial proteinase-K digestion experiment showed that the assemblies formed by both mutated proteins were resistant to protease activity, as indicated by the bands that were present after digestion in the Coomassie-stained polyacrylamide gels (S5 Fig). Differences were, however, noticeable after 20 minutes of digestion. Treatment with TasA D64A rendered resistance bands between 15 and 25 kDa and between 10 and 15 kDa, which progressively disappeared over time. A similar pattern was obtained for TasA K68A and D69A; however, the band between 10 and 15 kDa remained after 60 minutes of digestion, indicating that this variant is likely more resistant to digestion than TasA D64A.

**Fig 6.**
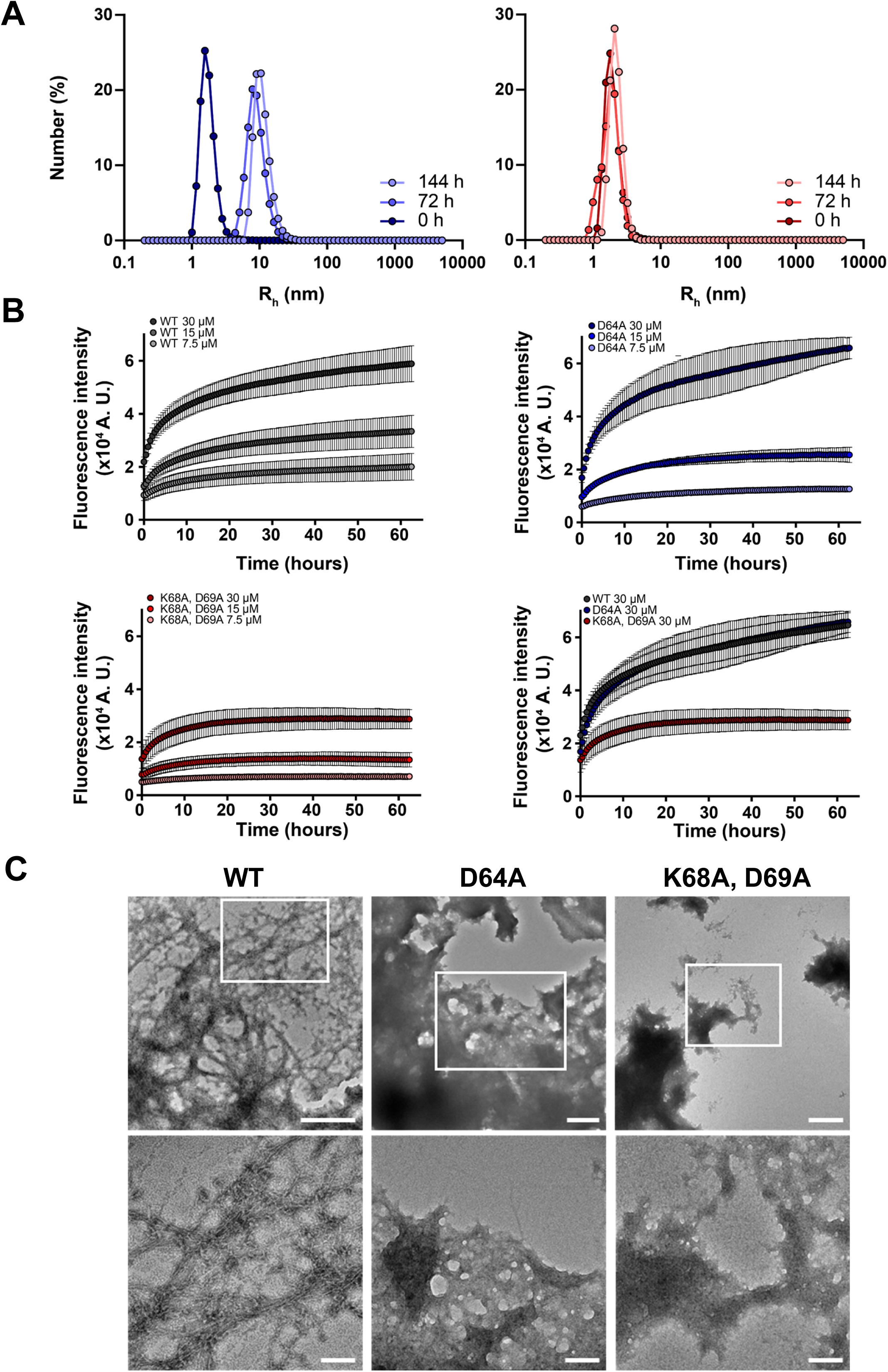
*In Vitro* Analysis of the TasA Variants Reveals Two Types of Molecular Behaviors. (A) Aggregation kinetics of the D69A allele (left) and K68A, D69A allele (right) as measured by DLS at different time points. (B) Kinetics of ThT fluorescence emission at 480 nm of the WT, D64A or K68A, D69A allele and superimposition of the plots corresponding to the 30 µM protein concentration. Experiments were performed at different protein concentrations. Average values of three independent experiments are represented. Error bars indicate the SEM. (C) Transmission electron microscopy micrographs of negatively stained samples of the WT and TasA variants. White squares indicate areas of the pictures that have been zoomed in. Scale bars = 500 and 100 nm from top to bottom.

All these results indicate that TasA D64A is a stable protein in cells with similar biochemical stability and behavior compared to TasA WT and is, in principle, not affected in its ability to form aggregates enriched in beta-sheets that are resistant to proteinase degradation. However, TasA K68A and D69A appear to be less stable than TasA WT and exhibit a stronger tendency to form aggregates with limited amyloid features. These two distinct molecular behaviors reflect the importance of these residues in the core’s contribution to TasA functionality and explain the two different phenotypes derived from these amino acid substitutions.

To further characterize the molecular conformation of TasA D64A, TasA K68A, and D69A amyloid fibrils, we employed solid-state NMR spectroscopy. Recombinant D64A or K68A, D69A mutant proteins were overexpressed in *E. coli* and ^13^C-labeled following protocols previously optimized ^10^. *In vitro* self-assembled filaments of the mutant proteins were investigated using magic-angle spinning SSNMR. We recorded a two-dimensional ^13^C-^13^C correlation experiment using a first cross-polarization ^1^H-^13^C step to reveal the conformation of rigid residues involved in the amyloid core. Comparison of ^13^C-^13^C spectral fingerprints (Fig 7A in red and green) showed that D64A, K68A, D69A assemblies exhibit broader NMR signals. ^13^C line-widths for the two mutants are ∼200-400 Hz (measured with full-width at half-height), suggesting a high propensity of the 2 mutants to adopt polymorphic amyloid structures. For WT assemblies (Fig 7A, in black), narrow peaks are observed with ^13^C line-widths of ∼100-200 Hz, indicating a low amount of structural polymorphism at the local level. These line-width values are not surprising for WT fibrils, since functional amyloids often lead to the aggregation of protein subunits into a unique polymorphic structure characterized by well-resolved SSNMR signals ^34^. The observation of much broader signals for the two mutants is nevertheless not unexpected for amyloid fibril samples, since pathological amyloids involved in misfolding diseases very often lead to broad NMR spectra due to the presence of severe local structural polymorphism ^35^. Next, we scrutinized the chemical shift values for the alanine Cα-Cβ spectral region. This spectral region is helpful for probing the secondary structure. Following the procedure described for WT TasA fibrils ^10^, we compared the two mutants to WT fibrils (Fig 7B). We observed two trends for the chemical shift values of the mutant fibril NMR signals: (i) several signals are conserved in their chemical shift values between the two mutants and the WT sample, suggesting that these residues have a conserved local conformation between the three samples. Nevertheless, (ii) numerous signals observed in the WT sample disappeared for the two mutants, indicating a partial loss of the structural integrity for the mutants. This observation was also made for other spectral regions of the two-dimensional experiments. Altogether, this chemical shift analysis suggests that the well-ordered amyloid core observed for WT TasA fibrils is only partially conserved for the two mutants, and additionally, a significant number of residues in the amyloid core of D64A, K68A, D69A mutants have lost their structural stability. Although the chemical shift analysis did not reveal a significant difference between D64A and K68A, D69A mutants, the comparison of cross-polarization (CP) and J-based (INEPT) polarization transfer showed a different behavior. While CP transfer reveals rigid residues of the structural core, INEPT signals probe the highly mobile residues of the assembly. We derived the CP/INEPT signal ratio from ^1^D ^13^C experiments (Fig 7C), and we observed that the ratio was comparable for WT and D64A samples (ratio ∼2) but decreased for K68A and D69A samples (ratio ∼1). The relative loss of CP signals for the K68A and D69A samples suggests overall more dynamic assemblies, in line with the TEM observations.

**Fig 7.**
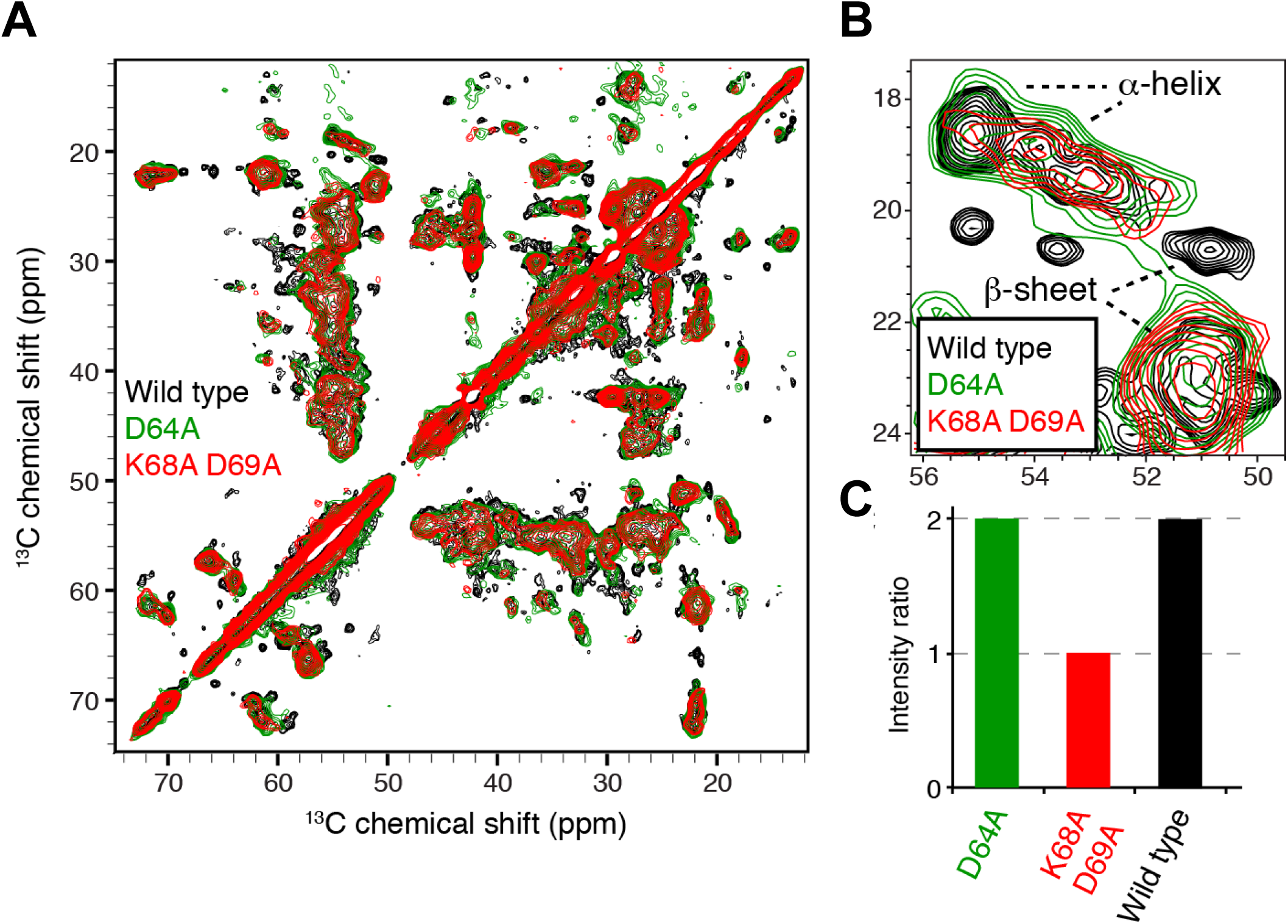
Recombinant WT TasA, TasA D64A or TasA K68A, D69A show Similar Structural Fingerprints. (A) 2D [^13^C]-[^13^C] SSNMR experiments of recombinant TasA WT (black), TasA D64A (green) and TasA K68A, D69A (red) filaments revealing the rigid residues. (B) Overlay of 2D SSNMR [^13^C]-[^13^C] spectra of TasA WT (black), TasA D64A (green) and TasA K68A and D69A (red) fibrils in the alanine Cα-Cβ spectral region. C) CP/INEPT signal ratio from ^1^D ^13^C experiments in samples of the WT TasA, TasA D64A and TasA K68A and D69A fibrils.

### *In vivo* characterization of the TasA variants demonstrates a complete loss of structural function in the K68A and D69A alleles

Previous experiments have demonstrated that the two amino acid substitutions introduced in the imperfect repeat region of AcTasA have an effect on morphology and biofilm formation and have revealed that i) the D64A allele shows a molecular behavior similar to that of the WT protein, which is able to self-assemble with a tendency to form dense aggregates, consisting of large bundles of fibers that exhibit amyloid properties and ii) the K68A, D69A protein is unstable, exhibits limited amyloid properties and a tendency toward the formation of aggregates instead of fibers.

Despite the loss of functionality of both TasA variants compared to the WT protein, the strains expressing these two versions of the protein still formed a pellicle, although it was different from that of the WT strain or the strain expressing the native protein. One possible explanation for this contradictory finding is the compensatory structural activity mediated by EPS, the other major component of the ECM in *B. subtilis* biofilms. To test this hypothesis, the EPS operon was deleted in all the strains expressing the different versions of TasA. The biofilm formation phenotype of the strains carrying the native or D64A variant proteins resembled that of the Δ*epsA-O* strain alone, in which a weak and fragmented pellicle floated in the liquid medium (Fig 8A). However, the phenotype of the strain expressing TasA_K68A, D69A_ mirrored that of the double Δ*tasA*, Δ*epsA-O* deletion strain, showing a complete absence of the pellicle (Fig 8A right image). According to these biofilm phenotypes, cells of the Δ*epsA-O*, Δ*eps-A-O* TasA_native_, and Δ*epsA-O* TasA_D64A_ strains were uniformly decorated with a network of fibers that was morphologically indistinguishable between these strains (Fig 8B). The identical growth dynamics of all the strains (Fig 8C) and the estimated cell density (Fig 8D) in MSgg led us to discard any growth defect of the strain expressing TasA_K68A, D69A_.

**Fig 8.**
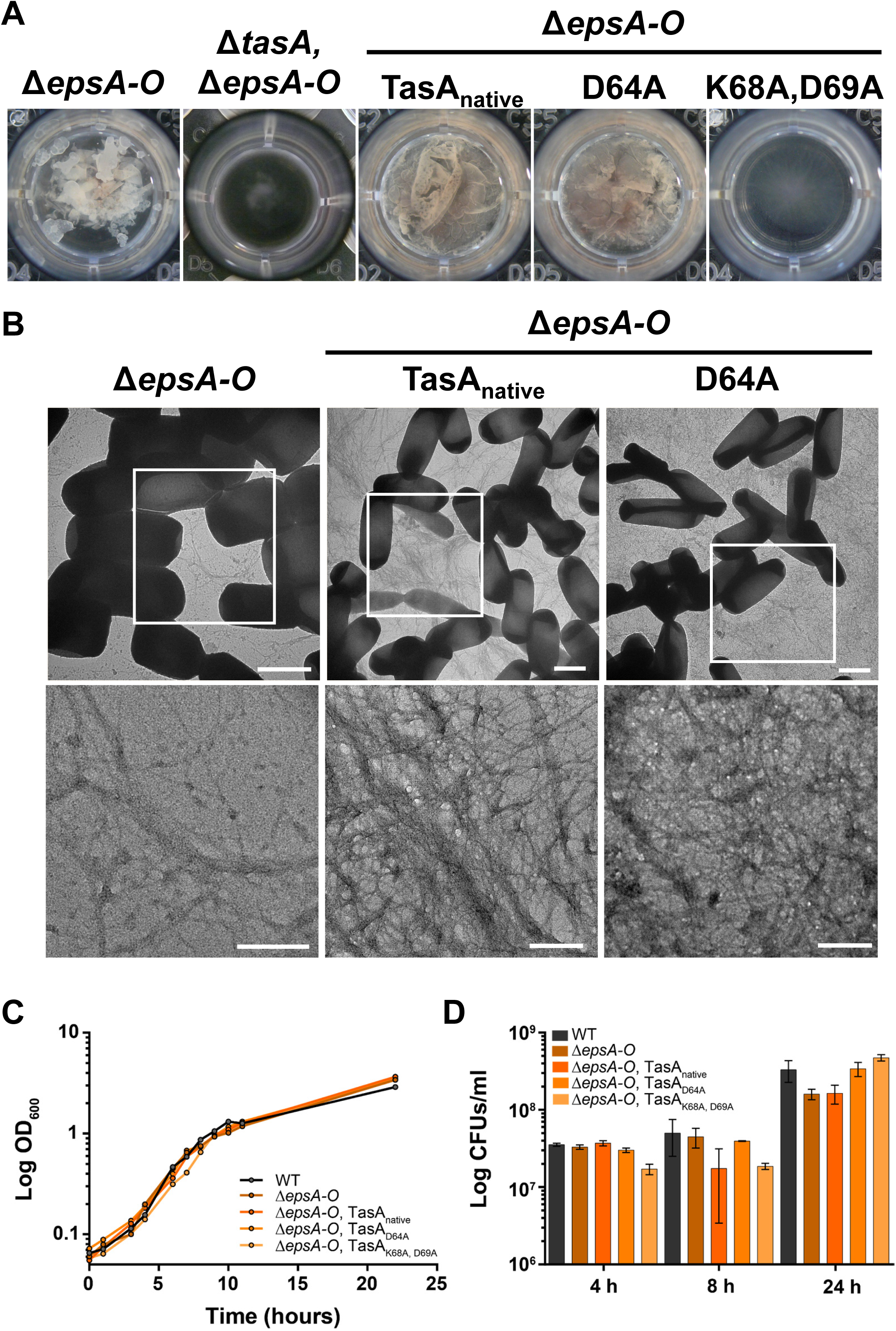
K68A and D69A Completely Abrogated Biofilm Formation. (A) The images show the biofilm formation phenotypes of the Δ*epsA-O*, Δ*tasA*Δ*epsA-O*, and Δ*epsA-O* strains carrying the different TasA variant alleles. (B) Transmission electron microscopy micrographs of negatively stained samples from the Δ*epsA-O* or Δ*epsA-O* strain carrying the native TasA allele or the D64A allele grown under biofilm-inducing conditions. White squares indicate areas of the images that have been zoomed in. Scale bars = 1 µm (top) and 200 nm (bottom). (C) Growth curves of the WT, Δ*epsA-O*, and Δ*epsA-O* strains carrying the different TasA variant alleles in liquid MSgg medium. (D) Colony counts at different time points of the growth curve of the strains WT, Δ*epsA-O*, and Δ*epsA-O* carrying the different TasA variant alleles in liquid MSgg medium.

As previously reported ^18^, the mixture of Δ*tasA* and Δ*eps* strains complement themselves when coinoculated in liquid medium, providing each other externally with their missing ECM components and restoring the wrinkled phenotype characteristic of *B. subtilis* biofilms. The same phenomenon takes place in solid medium, in which the coinoculation of both strains in a 1:1 proportion results in a phenotype very different from the other two mutant strains, leading to the formation of wrinkles (S6 Fig, top images). The three Δ*epsA-O* strains expressing the different versions of TasA showed a phenotype that resembled that of the Δ*eps* strain in solid medium (S6 Fig middle row images). Restoration of the wrinkly phenotype was only achieved when Δ*tasA* was cocultured with Δ*epsA-*O carrying the TasA_native_ or TasA_D64A_ alleles (S6 Fig, bottom images), demonstrating that the biofilm-defective phenotype of the Δ*epsA-O* TasA_K68A, D69A_ strain is mostly due to a lack of structural functionality of AcTasA and the inability of this strain to assemble a functional ECM.

## Discussion

The N-terminal domain of TasA contains two amyloid stretches and one sequence of imperfect amino acid repeats. Our study has proven that at least one of the predicted amyloidogenic stretches found in AcTasA, which exhibits amyloid properties, has the potential to contribute to the process of polymerization of TasA and that some highly conserved amino acids within this imperfect repeat are potential determinants of the amyloid nature and the related structural functionality of TasA. Unfortunately, we were unable to characterize the TasA variants lacking these amyloidogenic regions, as their deletion resulted in the degradation of the protein. These regions are known to be of utmost importance to explain the molecular behavior of amyloid proteins ^36, 37^, and given the ability of amyloids to self-assemble through protein‒protein interactions, these stretches serve novel approaches in synthetic biology and biotechnology, for example, to capture free amyloid peptides or proteins or even bacteria themselves when exhibiting these sequences on their surface ^38^. The fact that these mutated proteins are eliminated to undetectable levels and the different phenotypes of strains expressing their alleles compared to the Δ*tasA* strain or any other ECM-defective strain suggest a relevant contribution to the native folding features of TasA.

The unique amino acid substitution D64A in AcTasA was sufficient to render a mutated protein that retained amyloid features but was incapable of contributing to biofilm formation. The fact that this variant is able to complement biofilm formation, at least partially, when it is produced in a ΔEPS background and coinoculated with a Δ*tasA* strain, suggests that when both components are produced in the same cell, they are not completely functional. Indeed, this is reversed when they are expressed in different cell populations. We speculate that the fibers formed by TasA_D64A_, although functional from a structural and biochemical point of view, modify the surface of the cells, altering the deposition of the EPS in the ECM or disturbing any other putative interaction between the amyloid-like filaments and the EPS. Such a phenomenon would not be unprecedented, given that similar interactions between functional amyloids and polysaccharides have been reported in other bacterial species. In biofilms of *Streptomyces coelicolor,* the fimbriae that mediate adhesion to surfaces are composed of bundles of amyloid fibers formed by chaplins, functional amyloids involved in reducing the water surface tension and promoting the development of aerial structures ^39^. These fimbriae are maintained attached to the cell surface via their interaction with cellulose, which seems to serve as a template for the polymerization of the fimbriae ^40^. Moreover, a more recent work has proposed that coiled-coil domains in intermediate filament-like proteins have an intrinsic affinity for cellulose in different bacterial species ^41^, demonstrating a connection between specific domains in filamentous proteins and interactions with polysaccharides.

The substitutions K68A and D69A, which have been characterized in this work, were previously studied in which an additional function for TasA in cell physiology via its localization to the cell membrane was demonstrated ^19^. A strain expressing this version of TasA has a defective biofilm formation phenotype, which is disconnected from the cell membrane stability required for the proper physiological function of TasA ^19^. These amino acids are well conserved in the AcTasA region of imperfect repeats, and contrary to the other TasA variant investigated, their substitution affects the amyloid nature and stability of the entire protein, compromising its structural function in biofilm formation. Imperfect amino acid repeats are a common theme in many functional and nonfunctional amyloids ^21, 22, 25, 32, 42, 43^, a reason for their use for the localization of important residues for protein fibrillation. Indeed, the alteration of these regions can render altered amyloid behavior, either inhibiting or improving the process of amyloidogenesis.

A recent work has provided insight into the molecular structure of native TasA filaments purified from *B. subtilis* biofilms with atomic resolution ^16^. It was proposed that, as a monomer, the unstructured N-terminal end of the protein (amino acids from A28 to S41) along with the β1 sheet undergoes a conformational change that results in these residues extending away from the monomer folding into another beta-sheet that complements the next subunit in a process known as donor strand complementation and the refolding of β3. Based on this finding, the authors propose that the structural features of the native TasA filaments fit better in the category of amyloid-like proteins, as these fibers are neither a linear arrangement of globular subunits nor is their structure representative of the cross-β architecture typical of amyloid proteins. Nonetheless, it has been previously demonstrated that recombinant TasA *in vitro* shows the typical cross-β architecture in X-ray diffraction experiments, clearing any doubts regarding this question, at least concerning recombinant TasA. In addition, native filaments purified directly from *B. subtilis* biofilms, even if purified to homogeneity, may be affected by the presence of the accessory protein TapA in the cells ^18^; therefore, caution must be exercised. With the data that are available currently, the core region of TasA fits rather well in the proposed model in which the N-terminal domain of TasA is involved in donating the strand required for the assembly of the subunits and for the interactions that stabilize this strand. It is tempting to speculate that the studied amino acid substitutions in D64, K68, D69 participate in this process of stabilization of the donor strand; therefore, their variation, especially in K68A, D69A, interferes with this process, affecting the assembly of the filaments.

Taken together, our findings, along with those from previously published works, lead us to a better understanding of the polymerization process of TasA during biofilm formation and provide knowledge into the sequence determinants that promote the molecular behavior of functional amyloids.

## Materials and methods

### Bacterial strains and culture conditions

The bacterial strains used in this study are listed in S1 Table. Bacterial cultures were grown at 37 °C from frozen stocks on Luria-Bertani (LB: 1% tryptone (Oxoid), 0.5% yeast extract (Oxoid) and 0.5% NaCl) plates. Isolated bacteria were inoculated in appropriate medium. Biofilm assays were performed on MSgg medium: 100 mM morpholinepropane sulfonic acid (MOPS) (pH 7), 0.5% glycerol, 0.5% glutamate, 5 mM potassium phosphate (pH 7), 50 μg/ml tryptophan, 50 μg/ml phenylalanine, 50 μg/ml threonine, 2 mM MgCl_2_, 700 μM CaCl_2_, 50 μM FeCl_3_, 50 μM MnCl_2_, 2 μM thiamine, and 1 μM ZnCl_2_. For *in vitro* cell complementation assays with cell-free supernatants, the medium optimized for lipopeptide production (MOLP) was used and prepared as previously described. For cloning and plasmid replication, *Escherichia coli* DH5α was used. For protein purification, *Escherichia coli* BL21(DE3) was used. *Bacillus subtilis* 168 is a naturally competent domesticated strain used as a first step to transform the different constructs into *Bacillus subtilis* NCIB3610 by SPP1 phage-mediated generalized transduction. The final antibiotic concentrations for *B. subtilis* were MLS (1 μg/ml erythromycin, 25 μg/ml lincomycin), spectinomycin (100 μg/ml), tetracycline (10 μg/ml), chloramphenicol (5 μg/ml), and kanamycin (10 μg/ml). For the selection of plasmids in *E. coli*, ampicillin at 100 μg/ml was used.

### Plasmid and strain construction

TasA was purified using the pDFR6 (pET22b-*tasA*) plasmid containing the ORF of TasA without the signal peptide or the stop codon, which was constructed as previously described. To purify TasA_D64A_ and TasA_K68A, D69A_, the ORF of the mutated alleles excluding the signal peptide were amplified from strains JC78 and JC81 using the primers TasA_Exp_C_NdeI_F and TasA_Exp_C_XhoI_R. The resulting PCR products were then cloned into pET22b by digestion with NdeI and XhoI, and the final plasmids were maintained in *E. coli* (strains JC104 and JC106, respectively).

The region corresponding to the amyloid core of TasA was amplified from the *tasA* ORF using primers TasA_Exp_C_NdeI_F and TasA_Exp_C_XhoI_R. The resulting PCR product was digested using NdeI and XhoI and cloned into pET22b. The final plasmid was maintained in *E. coli* (strain JC118).

Strains JC72, JC75, JC76, JC80 and JC82 were generated by site-directed mutagenesis using the primer pairs del82-88, del82-88-antisense, del108-116, del108-116-antisense KD_AA_35-36, KD_AA_35-36_antisense, F_A_72, F_A_72_antisense, E_A_82, E_A_82_antisense, G_A_96, G_A_96_antisense and D_A_64, D_A_64_antisense with a QuickChange Lightning Site Directed Mutagenesis Kit (Agilent Technologies), following the manufacturer’s instructions.

Strains JC209, JC222 and JC224 containing the mCherry fusions of the native protein or the TasA_K68A, D69A_ or TasA_D64A_ variants, respectively, were generated using the oligos op-tasA_fw, op-tasA_rv, mCherry_fw, mCherry_rv, tasA_downstream_fw, and tasA_downstream_rv and the NEBuilder HiFi DNA Assembly Cloning Kit (New England Biolabs).

All of the *B. subtilis* strains generated were constructed by transforming *B. subtilis* 168 via its natural competence and then using the positive clones as donors for transferring the constructs into *B. subtilis* NCIB3610 via generalized SPP1 phage transduction ^44^

### Bioinformatic analysis of the TasA sequence

For the prediction of amyloidogenic regions within the N-terminal region of the TasA protein sequence, we used different sequence-based tools that are freely available online and applied different computational methods ^45^. MetAmyl ^46^ and AmylPred 2.0 ^47^ were used as consensus predictors, as both applications integrate the results provided by different online tools that predict protein aggregation or amyloidogenicity based on sequence analysis. APPNN ^48^ and FISH amyloid ^49^ were used as predictors based on machine-learning approaches. The consensus regions obtained from the different methods were selected for further experimental analyses.

The analysis of imperfect amino acid repeats present within the TasA sequence was performed using the webtool RADAR ^50^, available at the EMBL-EBI website.

The structure prediction of WT TasA, TasA_D64A_ and TasA_K68A, D69A_ was performed using the crystal structure of the TasA monomer as a template. Structures were predicted using AlphaFold ^51^ in a Google Colab notebook (ColabFold) ^52^. Protein models were visualized and analyzed using UCSF ChimeraX (v1.3) ^53^.

### Biofilm formation assays

To analyze biofilm formation and the colony architecture on plates and pellicle formation in liquid medium, the different strains were grown in LB plates at 37 °C overnight. Then, suspensions of the corresponding strains were made in PBS, and these were adjusted to an OD_600_ of 1. To analyze biofilm formation on the plates, 2 µl drops of the bacterial suspensions were spotted onto the surface of MSgg plates that were incubated at 30 °C. Pictures were taken at 24, 48 or 72 hours to observe the colony architecture at different time points. To analyze the formation of pellicles in the air liquid interface of liquid media, 10 µl of each bacterial suspension was added to 1 ml of liquid MSgg medium in 24-well plates. Then, the plates were incubated at 30 °C without agitation or aeration for 48 hours.

### Biofilm fractionation

The presence of TasA was analyzed in the studied strains by fractioning the biofilms in medium, cells or ECM as previously described ^9^. To have sufficient biological material, one 24-well plate was used for each bacterial strain, of which only the 8 central wells were used. Briefly, pellicles were carefully lifted from the wells and placed in a separate tube containing 8 ml of MS medium (MSgg without glycerol and glutamate). The remaining spent medium of the well was filtered sterilized and kept at 4 °C for further analysis. Then, to separate the cells from the ECM, the pellicles were subjected to mild sonication (approximately 10 pulses of 30 s in a Branson 450 digital sonifier, with 30 s pauses in ice), and the resulting suspensions were centrifuged at 9000 x g to separate the ECM from the cells. The supernatant (ECM fraction) was filtered, sterilized and kept at 4 °C. The cell fraction was resuspended in 8 ml of MS medium and kept at 4 °C until further processing.

### Synthetic peptides

Peptides corresponding to the amyloidogenic regions detected within the TasA amyloid core were synthetically produced and purified by Proteogenix (Schiltigheim, France). The sequence of the peptides was: for peptide LG-13 NH_2_-LAIKEVMALNYG-COOH and for peptide DG-14 NH_2_-DFLSQFEVTLLTVG-COOH. A stock solution of each peptide was prepared in DMSO. Then, the peptides were diluted in Tris 20 mM and NaCl 50 mM to prepare the working solution for the different experiments at the appropriate concentration.

### Protein precipitation

To precipitate the proteins of each biofilm fraction, 2 ml of the medium and ECM fractions were directly treated with 10% trichloroacetic acid (TCA) and incubated on ice for 1 h. For the cell fractions, 2 ml of the suspension was treated with 0.1 mg/ml lysozyme and incubated at 37 °C for 30 min. Then, the lysozyme-treated suspension was subjected to TCA treatment as described above. After incubation, proteins were collected by centrifugation at 13000 x g for 20 min at 4 °C. The pellets were washed twice with cold acetone and dried by air. The proteins were resolubilized in 8 M urea, 100 mM Tris, and 150 mM NaCl by incubating at 100 °C for 5 mins.

### Protein expression and purification

Protein was expressed and purified as previously described ^10^. Briefly, *E. coli* BL21(DE3) cells were freshly transformed with the pET22b plasmid containing either WT TasA (pDFR6 plasmid), the different mutated versions of the protein (D64A, and K68A, D69A) or the amyloid core region. Colonies were selected from the plates, resuspended in 10 mL of LB with 100 µg/mL ampicillin and incubated overnight at 37 °C with shaking. This preinoculum was then used to inoculate 500 mL of LB + ampicillin, and the culture was incubated at 37 °C until an OD600 of 0.7–0.8 was reached. Next, the culture was induced with 1 mM isopropyl β-D-1-thiogalactopyranoside (IPTG) and incubated O/N at 30 °C with shaking to induce the formation of inclusion bodies. After that, cells were harvested by centrifugation (5000 × g, 15 min, 4 °C), resuspended in buffer A (Tris 50 mM, 150 mM NaCl, pH 8), and then centrifuged again. These pellets were stored frozen at −80 °C until use. After thawing, cells were resuspended in buffer A and broken down by sonication on ice using a Branson 450 digital sonifier (3 × 45 s, 60% amplitude). After sonication, the lysates were centrifuged (15,000 × g, 60 min, 4 °C), and the supernatant was discarded, as proteins were mainly expressed in inclusion bodies. The proteinaceous pellet was resuspended in buffer A supplemented with 2% Triton X-100, incubated at 37 °C with shaking for 20 min to further eliminate any remaining cell debris, and centrifuged (15,000 × g, 10 min, 4 °C). The pellet was then extensively washed with buffer A (37 °C, 2 h), centrifuged (15,000 × g for 10 min, 4 °C), resuspended in denaturing buffer (Tris 50 mM NaCl 500 mM, 6 M GuHCl), and incubated at 60 °C overnight to completely solubilize the inclusion bodies. Lysates were clarified via sonication on ice (3 × 45 s, 60% amplitude) and centrifugation (15,000 × g, 1 h, 16 °C) and were then passed through a 0.45-µm filter prior to affinity chromatography. Proteins were purified using an AKTA Start FPLC system (GE Healthcare). The lysates were loaded into a HisTrap HP 5 mL column (GE Healthcare) previously equilibrated with binding buffer (50 mM Tris, 0.5 M NaCl, 20 mM imidazole, 8 M urea, pH 8). Protein was eluted from the column with elution buffer (50 mM Tris, 0.5 M NaCl, 500 mM imidazole, 8 M urea, pH 8). After the affinity chromatography step, the buffer was exchanged with 1% acetic acid at pH 3 and 0.02% sodium azide by using a HiPrep 26/10 desalting column (GE Healthcare). This ensured that the proteins were maintained in their monomeric form. The purified proteins were stored under these conditions at 4 °C (maximum 1 month) until further use.

### Assembly of TasA filaments

To assemble the TasA filaments *in vitro* from the corresponding monomeric purified proteins, first, the acidic buffer was removed and exchanged with Tris 20 mM NaCl 50 mM using FPLC and a HiPrep 26/10 desalting column as described above. Filaments were assembled with a protein concentration above 1 mg/ml and incubated at 30 °C in standing conditions for one week.

### SDS‒PAGE and Western blot

Protein samples from the different assays were diluted in 2x Laemmli sample buffer (Bio-Rad) and heated at 100 °C for 5 min. Proteins were separated by SDS‒PAGE in 12% acrylamide gels and then transferred onto PVDF membranes using a Trans-Blot Turbo Transfer System (Bio-Rad) and PVDF transfer packs (Bio-Rad). For immunodetection of TasA, the membranes were incubated with an anti-TasA antibody (rabbit) at a 1:20,000 dilution in Pierce Protein-Free (TBS) blocking buffer (Thermo Fisher). A secondary anti-rabbit IgG antibody conjugated to horseradish peroxidase (Bio-Rad) was used at a 1:3000 dilution in the same buffer. The membranes were developed using Pierce ECL Western blotting Substrate (Thermo Fisher).

### Transmission electron microscopy

To visualize the fibers formed by the different TasA variants, the assembled protein samples were deposited over carbon-coated copper grids and incubated for 1 hour. Next, the excess sample was bloated off the grid, and the samples were negatively stained by floating the grids in a 2% uranyl acetate solution for 30 s and then washed by floating the grid in distilled water for 30 seconds.

To visualize the filaments formed by the synthetic peptides corresponding to the amyloidogenic regions of the amyloid core of TasA, the samples were taken from the microplates used in the thioflavin-T binding experiments after saturation of the signal (36 h of incubation) and were deposited in grids and negatively stained as described above.

To study the morphology of fibers on the surface of cells, pellicles of the different strains were grown as described above. After 48 h of incubation, carbon-coated copper grids were deposited into the wells over the fully formed pellicles and incubated for 2 h at 30 °C in standing conditions. After incubation, the grids were washed twice with distilled water for 30 s, negatively stained with 1% uranyl acetate for 20 s and washed again once with distilled water for 30 s.

All the samples were visualized in an FEI Tecnai G2 20 TWIN transmission electron microscope at an accelerating voltage of 80 kV. The images were taken using a side-mounted CCD Olympus Veleta with 2k x 2k Mp.

### Confocal laser scanning microscopy

The localization of TasA was studied using a TasA-mCherry translational fusion in a Δ*tasA* genetic background (see S1 Table). Strains JC209, JC222 and JC224 containing the WT protein fused to mCherry or the fusion of the different variants, respectively, were grown in solid medium under biofilm-inducing conditions, as described above, and the different colonies were taken at 48 h and resuspended in distilled water using a 25 5/8 G needle. Bacterial suspensions were treated with CellBrite Fix 488 Membrane Stain (Biotium, stock solution at 1000X) at a final concentration of 1X. Images of the stained bacteria were acquired sequentially to obtain images from the membrane stain and mCherry fluorescence. The CellBrite image was acquired by exciting the dye at 488 nm and recording the emissions from 498 to 553 nm, followed by a second acquisition corresponding to the mCherry fluorescence with excitation at 561 nm and recording of the emissions from 572 to 665 nm.

All images were obtained by visualizing the samples using an inverted Leica SP5 system with a 63x NA 1.4 HCX PL APO oil-immersion objective. For each experiment, the laser settings, scan speed, PMT or HyD detector gain, and pinhole aperture were kept constant for all of the acquired images.

### Thioflavin-T binding assays

To ensure that the assay is performed when TasA is in monomeric form, the purified proteins or the amyloid core were buffer exchanged and immediately used afterward in 20 mM Tris and 50 mM NaCl. The assay was performed in 96-well microplates. TasA or the different variants were mixed at different final concentrations of 7.5, 15 or 30 µM with 30 µM thioflavin-T in a final volume of 200 µl. In the experiments with the amyloid core region, the purified core was mixed at a final concentration of 15, 30 or 60 µM with thioflavin-T under the same conditions. Measurements were performed at 30 °C every 30 min in a Fluostar Omega plate reader (BMG Labtech) equipped with 450 nm and 480 nm filters for excitation and emission, respectively.

For the experiments performed with peptide samples, working solutions of both peptides were prepared in 20 mM Tris and 50 mM NaCl, and the peptides were used at a final concentration of 200 µM.

### Dynamic light scattering experiments

Freshly purified TasA, TasA amyloid core or the different TasA variants were buffer exchanged to 50 mM Tris and 20 mM NaCl containing 0.02% sodium azide immediately prior to the experiment. The final concentration was adjusted to 30 µM for all the samples, and these were incubated at 30 °C in standing conditions. Samples were taken at different time points to measure particle size in a Zetasizer Nano ZS (Malvern) equipped with a 632.8 nm laser as the excitation source and using 1 cm path length polystyrene cuvettes. Zetasizer software version 8.01 (Malvern) was used for data analysis.

### Circular dichroism measurements

WT TasA or the different variants were freshly purified and buffer exchanged to 20 mM phosphate buffer at pH 7 containing 0.02% sodium azide. For the thermal denaturation experiments, changes in ellipticity at 222 nm were recorded as a function of the increasing temperature. The thermal denaturation spectra were recorded by measuring the spectra of all the samples in 1 mm quartz cuvettes between 210 and 230 nm, with temperature steps of 10 °C and incubation times of 5 minutes between each step. Measurements were performed with the following parameters: resolution of 1 nm, bandwidth of 1 nm, sensitivity of 100 mdeg and 5 accumulations.

### SSNMR spectroscopy

All experiments were recorded at 600 MHz on a Bruker spectrometer equipped with a triple resonance 3.2 mm probe using a magic-angle spinning frequency of 11 kHz. The sample temperature was set to 278 K, and chemical shifts were calibrated using DSS as an internal reference. A decoupling strength of 90 kHz was used with small phase incremental alternation with 64 steps (SPINAL-64). 1D ^13^C experiments on LG-13 and DG-14 were recorded using a cross-polarization contact time of 1 ms and 20k scans. 2D ^13^C-^13^C experiments were recorded using a proton-driven spin diffusion mixing time of 50 ms.

### Partial proteinase-K digestion experiments

Experiments were performed as previously described ^10, 54^. Briefly, 50 µg of assembled WT TasA, the different variants or the amyloid core were incubated for 1 h at 37 °C with 0.02 mg/ml proteinase K in 20 mM Tris pH 8 and 50 mM NaCl, and samples were taken at different time points for further analysis (1, 5, 15, 20, 30, 45 or 60 min). Reactions were stopped in each sample by the addition of 1 vol. Laemmli buffer and heated at 100 °C for 5 min. Then, samples were separated by SDS‒PAGE and stained with Coomassie Brilliant Blue.

## Acknowledgments

We thank Juan Félix López Téllez from the Nanoimaging Unit of the Plataforma Bionand, Instituto de Investigación Biomédica de Málaga (IBIMA), for his technical assistance in the transmission electron microscopy imaging and analysis. We thank José Luis Zafra Paredes and José María Montenegro Martos from the Vibrational Spectroscopy unit and the Electronic Spectroscopy unit, respectively, of the Servicios Centrales de Apoyo a la Investigación (SCAI) from the University of Málaga for their technical assistance with the circular dichroism and dynamic light scattering experiments. We thank David Vela Corcía for his help with the protein modelling and hydrogen bond analysis. We also thank Estelle Morvan, Axelle Grélard and the Biophysical and Structural Chemistry Platform at IECB, CNRS UAR 3033, INSERM US001, University of Bordeaux.

## Author contributions

D. Romero conceived the study. D. Romero, and J. Cámara-Almirón. designed the experiments. J. Cámara-Almirón performed the main experimental work. L. Domínguez-García. performed the experiments with the AcTasA and provided assistance with the experimental work. A. Loquet and B. Habenstein designed the SSNMR spectroscopy experiments. N. El Mammeri and A. Lends carried out the SSNMR spectroscopy experiments. J. Cámara-Almirón, D. Romero, A. Loquet and N. El Mammeri wrote the manuscript. L. Domínguez-García, N. El Mammeri, A. Loquet, B. Habenstein, A. Lends, and A. de Vicente contributed critically to writing the final version of the manuscript.

## Supporting information

**S1 Fig. All the TasA Variants Exhibit Normal Localization in the Cell Membrane.** Confocal microscopy micrographs of cells carrying the different TasA alleles fused to a mCherry reporter and stained with a membrane dye (CellBrite). Scale bars = 5 µm.

**S2 Fig. Transmission Electron Microscopy Images of Cells Carrying the Native Proteins or the Protein Variants.** Transmission electron microscopy images of negatively stained samples from cells carrying the native or the D64A or K69 and D69A alleles or Δ*tasA* cells growing under biofilm inducing conditions in MSgg medium. White squares indicate areas of the images that have been zoomed in. Scale bars = 500 nm (left), 200 nm (middle) and 100 nm (right).

**S3 Fig. Structure Comparison between the Crystal Structure of the TasA Monomer and the Predicted Model.** Structure comparison between the crystal structure of TasA and the model predicted by AlphaFold. The numbering in the scheme follows the representation described for the crystal structure of TasA in its monomeric form published by Diehl et al. ^11^. The image on the right shows the superimposition of the two structures. The coloring scheme indicates the position of the residue in the sequence of the protein, where blue indicates the N-terminus and red indicates the C-terminus. Warmer colors indicate proximity to the C-terminal end.

**S4 Fig. The TasA Variants Show Differences in Thermal Stability Compared to the WT Protein.** Mean residue ellipticity at 222 nm measured as a function of temperature in the WT or the different TasA variants.

**S5 Fig. Comassie Stained SDS‒PAGE Gel of the D64A (left) or K68A and D69A Samples Digested with Proteinase K at Different Time Points.** The M lane indicates the molecular marker. Each lane contains a sample corresponding to a specific digestion time. SDS‒PAGE gel images have been cropped and spliced for illustrative purposes. The dashed lines over the gel images indicate the boundaries of the image splicing. The two slices in the two different gel images are derived from a single gel.

**S6 Fig. The TasA D64A Allele, but not the K68A or D69A Allele, is Able to Perform Extracellular Complementation When Mixed with a Δ*tasA* Strain**. Colony morphology phenotypes of Δ*tasA* and Δ*epsA-O* strains, of the coinoculation between Δ*tasA* and Δ*epsA-O*, of the Δ*epsA-O* strains carrying the different TasA alleles and of the coinoculation of these strains with the Δ*tasA* strain. Scale bars = 1 cm.

