## Supporting_information for "Molecular characterization of the N-terminal half of TasA during functional amyloid assembly and its contribution to *Bacillus subtilis* biofilm formation"

**-Supplementary figures 1 to 6**

**-Supplementary tables 1 and 2**

**S1 Fig.**

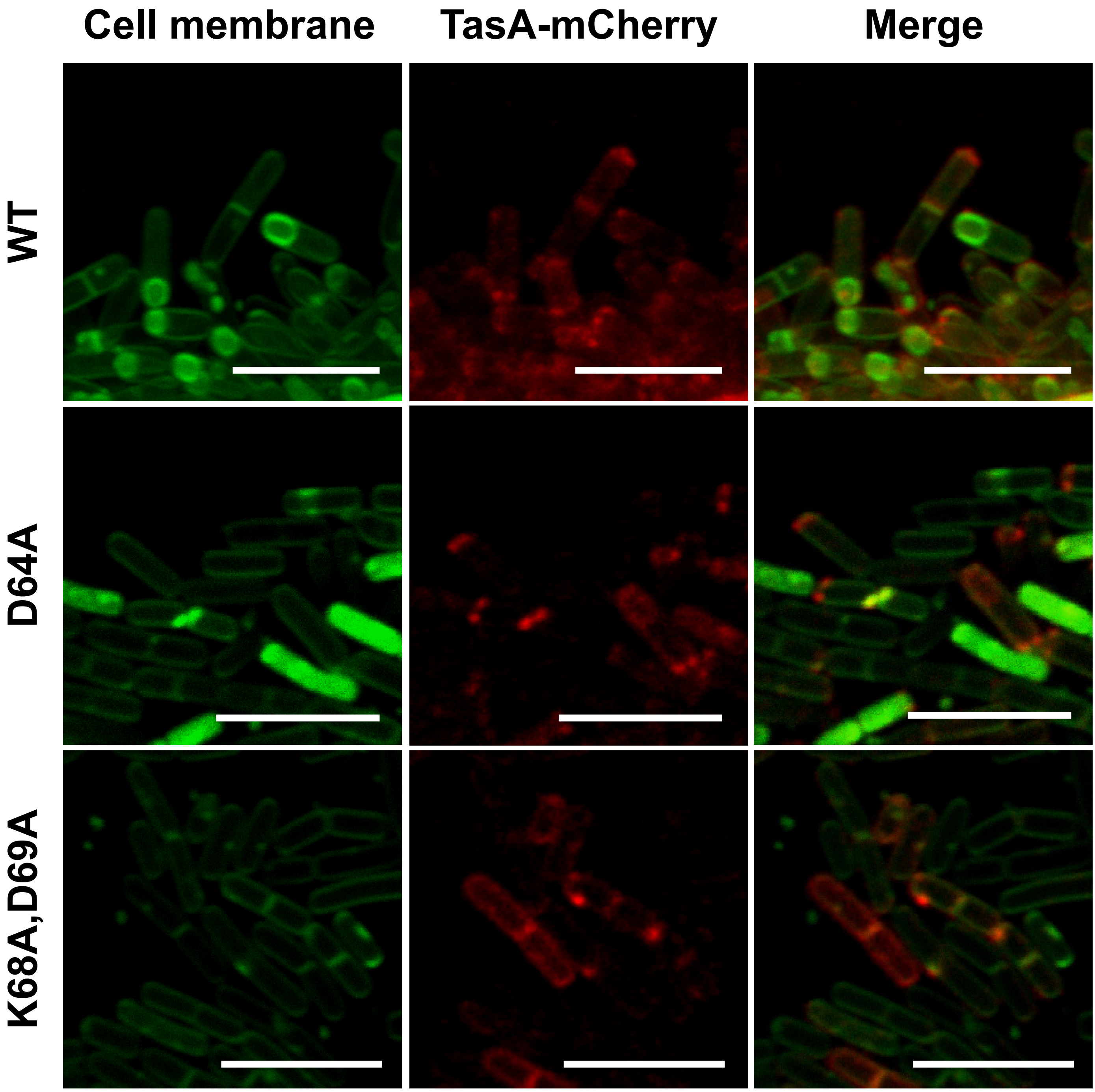

**S2 Fig.**

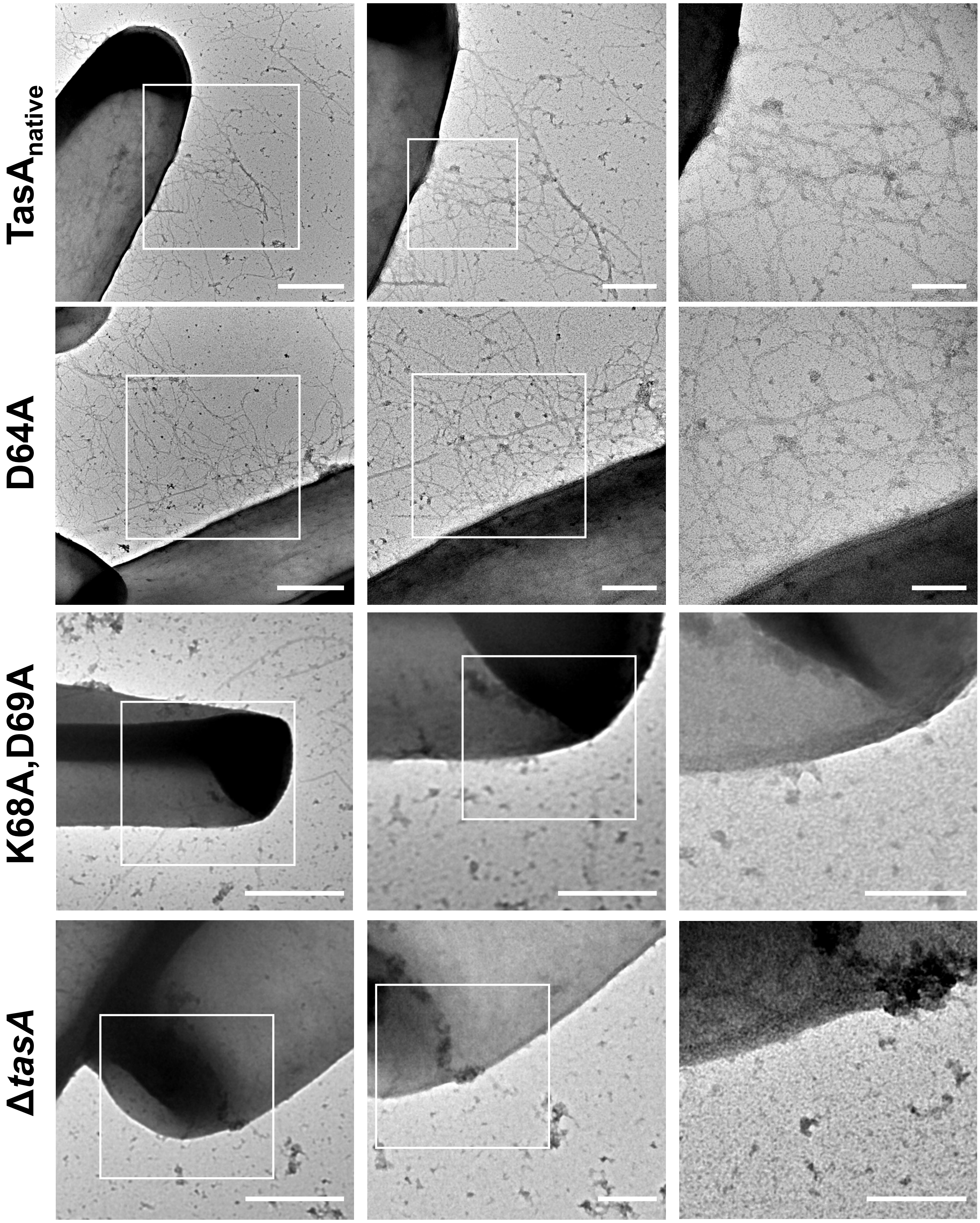

**S3 Fig.**

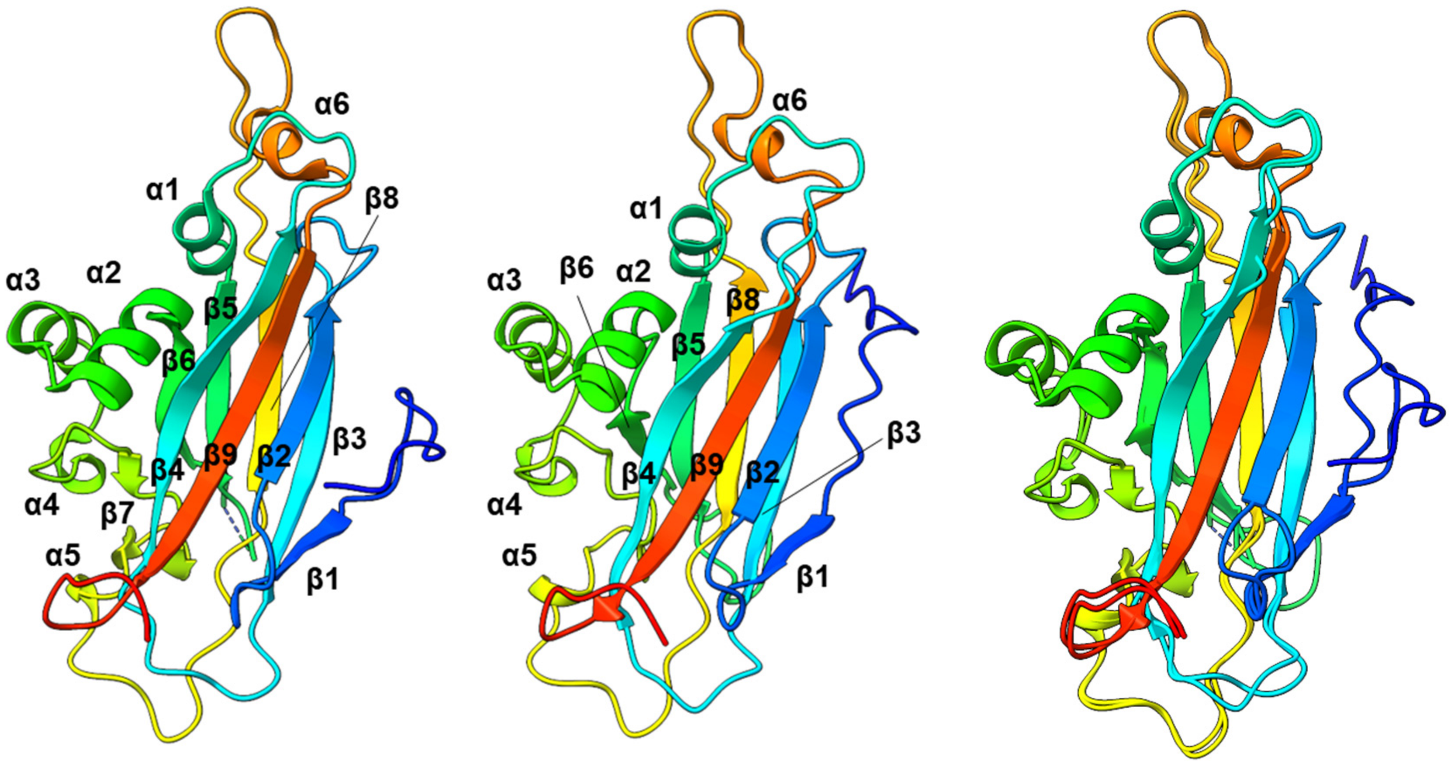


5OF1
(TasA determined
crystal structure)

Model of TasA
predicted by
AlphaFold

Structure alignment

**S4 Fig.**

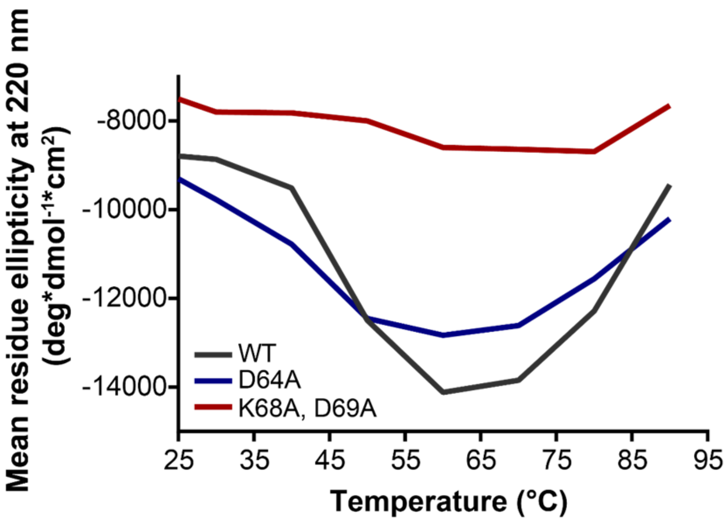

Mean residue ellipticity at 220 nm
(deg*dmol-1*cm2)
-8000
-10000
-12000
-14000
WT
D64A
K68A, D69A
25
35
45
55
65
75
85
95
Temperature (°C)

### S5 Fig.

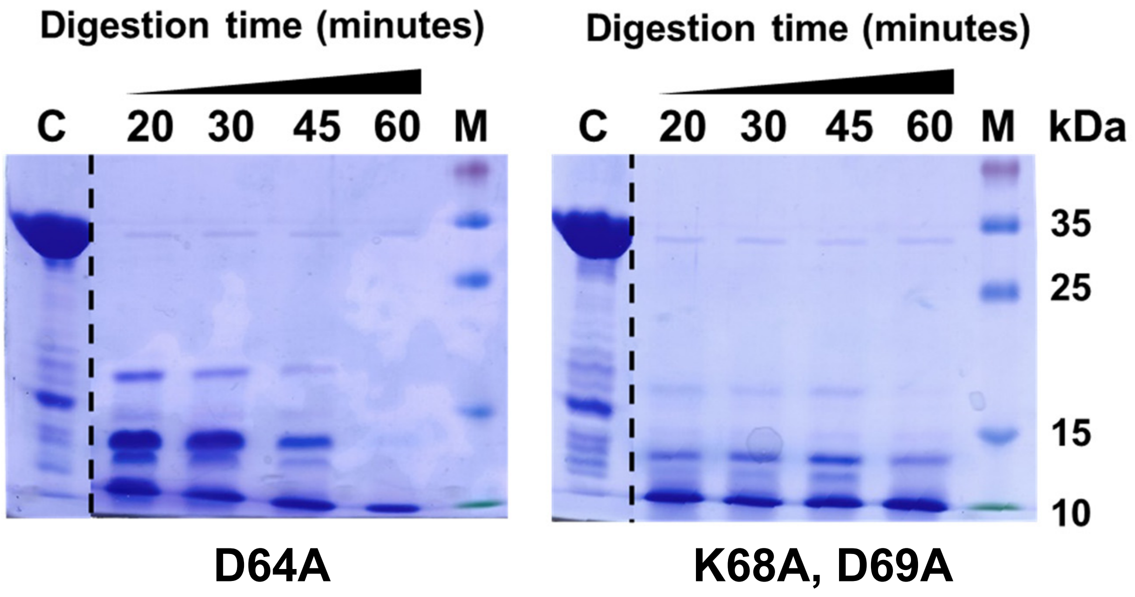

Digestion time (minutes)
C
20
30
45
60
M
D64A
Digestion time (minutes)
C
20
30
45
60
M
kDa
35
25
15
10
K68A, D69A

**S6 Fig.**

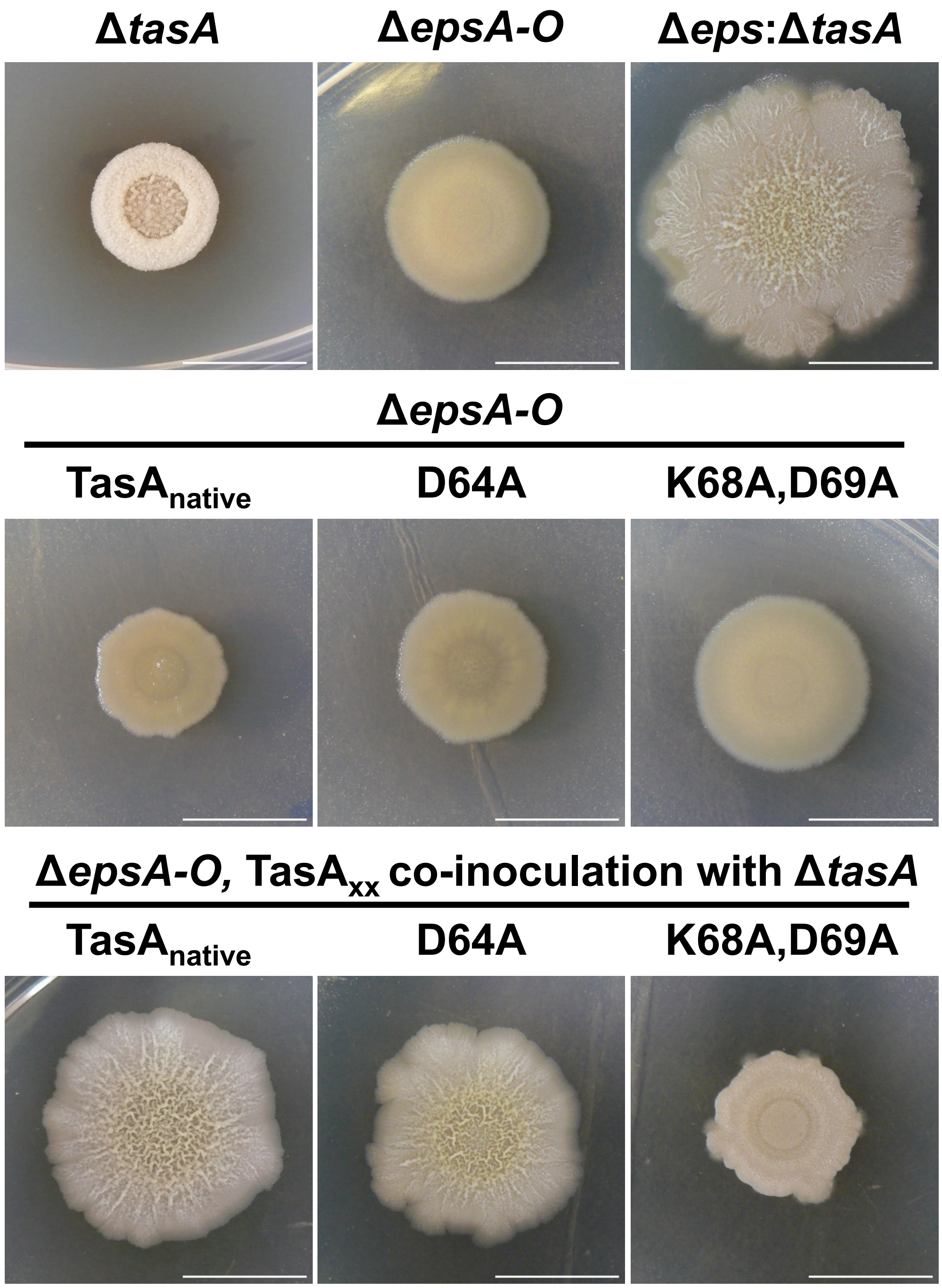
ΔtasA
ΔepsA-O
Δeps:ΔtasA
ΔepsA-O
TasAnative
D64A
K68A,D69A
ΔepsA-O, TasAxx co-inoculation with ΔtasA
TasAnative
D64A
K68A,D69A

**S1 Table**

Bacterial strains used in this study

| Bacterial strain | Genotype | Source |
|---|---|---|
| *Bacillus subtilis* 168 | Laboratory strain | Laboratory collection |
| *Bacillus subtilis* NCIB3610 | Wild type. Undomesticated strain | Laboratory collection |
| CA017 | *Bacillus subtilis* NCIB3610 *tasA::km* | (Romero *et al*., 2010) (13) |
| SSB488 | *Bacillus subtilis* NCIB3610<br>*epsA-O::tet* | (Branda *et al*., 2006) (10) |
| SSB149 | *Bacillus subtilis* NCIB3610<br>*(tapA-sipW-tasA)::spc* | (Branda *et al*., 2006) (10) |
| JC209 | *Bacillus subtilis* NCIB3610<br>*tasA::km*<br>*amyE::(tapA-sipW-tasA-mCherry) (mls)* | This work |
| JC70 | *Bacillus subtilis* NCIB3610<br>*(tapA-sipW-tasA)::spc*<br>*lacA::(tapA-sipW-tasA*$_{native}$*) (mls)* | (Cámara-Almirón *et al*., 2020) (19) |
| JC81 | *Bacillus subtilis* NCIB3610<br>*(tapA-sipW-tasA)::spc*<br>*lacA::(tapA-sipW-tasA*$_{(Lys68Ala,\ Asp69Ala)}$*) (mls)* | (Cámara-Almirón *et al*., 2020) (19) |
| JC78 | *Bacillus subtilis* NCIB3610<br>*(tapA-sipW-tasA)::spc*<br>*lacA::(tapA-sipW-tasA*$_{Asp64Ala}$*) (mls)* | This work |
| JC72 | *Bacillus subtilis* NCIB3610<br>*(tapA-sipW-tasA)::spc*<br>*lacA::(tapA-sipW-tasA*$_{\Delta 82-88}$*) (mls)* | This work |
| JC75 | *Bacillus subtilis* NCIB3610<br>*(tapA-sipW-tasA)::spc*<br>*lacA::(tapA-sipW-tasA*$_{\Delta 108-116}$*) (mls)* | This work |
| JC76 | *Bacillus subtilis* NCIB3610<br>*(tapA-sipW-tasA)::spc*<br>*lacA::(tapA-sipW-tasA*$_{Glu82Ala}$*) (mls)* | This work |
| JC77 | *Bacillus subtilis* NCIB3610<br>*(tapA-sipW-tasA)::spc*<br>*lacA::(tapA-sipW-tasA*$_{Phe72Ala}$*) (mls)* | This work |
| JC80 | *Bacillus subtilis* NCIB3610<br>*(tapA-sipW-tasA)::spc*<br>*lacA::(tapA-sipW-tasA*$_{Lys35Ala,\ Asp36Ala}$*) (mls)* | This work |
| JC82 | *Bacillus subtilis* NCIB3610<br>*(tapA-sipW-tasA)::spc*<br>*lacA::(tapA-sipW-tasA*$_{Gly96Ala}$*) (mls)* | This work |
| JC222 | *Bacillus subtilis* NCIB3610<br>*tasA::km*<br>*amyE::(tapA-sipW-tasA*$_{Lys68Ala,\ Asp69Ala}$*-mCherry) (mls)* | This work |
| JC224 | *Bacillus subtilis* NCIB3610<br>*tasA::km*<br>*amyE::(tapA-sipW-tasA*$_{Asp64Ala}$*-mCherry) (mls)* | This work |
| JC226 | *Bacillus subtilis* NCIB3610<br>*(tapA-sipW-tasA)::spc*<br>*lacA::(tapA-sipW-tasA*$_{native}$*) (mls)*<br>*epsA-O::tet* | This work |

| | | |
|---|---|---|
| JC228 | *Bacillus subtilis* NCIB3610<br>*(tapA-sipW-tasA)::spc*<br>*lacA::(tapA-sipW-tasA*$_{Asp64Ala}$*) (mls)*<br>*epsA-O::tet* | This work |
| JC231 | *Bacillus subtilis* NCIB3610<br>*(tapA-sipW-tasA)::spc*<br>*lacA::(tapA-sipW-tasA*$_{(Lys68Ala,\ Asp69Ala)}$*) (mls)*<br>*epsA-O::tet* | This work |
| JC118 | *E. coli* DH5α<br>*pET22b-TasA*$_{K35-K144}$ *(AcTasA)* | This work |
| JC104 | *E. coli* DH5α<br>*pET22b-TasA*$_{D64A}$ | This work |
| JC106 | *E. coli* DH5α<br>*pET22b-TasA*$_{K68A,\ D69A}$ | This work |

**S2 Table**

Primers used in this study

| Name | Sequence (5' – 3') | Purpose |
|---|---|---|
| Am_core_exp_C_Rv | aaaaactcgagtttagcagacatcaaatacaagtc | Cloning of the amyloid core of TasA into pET22b |
| Am_core_exp_C_Fw | aaaaacatatgaaggatgctacttttgcatcagg | Cloning of the amyloid core of TasA into pET22b |
| TasA_Exp_C_NdeI_F | aaaaacatatggcatttaacgacattaaatcaaa | Cloning of TasA D64A and TasA K68A, D69A into pET22b |
| TasA_Exp_C_XhoI_R | aaaaactcgagattttatcctcgctatgcgcttt | Cloning of TasA D64A and TasA K68A, D69A into pET22b |
| del82-88 | caatttgaaaataacggatcacttgcgatcaaatatggagattttaaagcaaac | Deletion of amino acids 82-86 from the TasA sequence by site-directed mutagenesis |
| del82-88-antisense | gtttgctttaaaatctccatatttgatcgcaagtgatccgttattttcaaattg | Deletion of amino acids 82-86 from the TasA sequence by site-directed mutagenesis |
| del108-116 | cagaagatttcctcagcggaaaagagggcggcaatg | Deletion of amino acids 108-116 from the TasA sequence by site-directed mutagenesis |
| del108-116-antisense | cattgccgccctcttttccgctgaggaaatcttctg | Deletion of amino acids 108-116 from the TasA sequence by site-directed mutagenesis |
| KD_AA_35-36 | ctgatgcaaaagtagcagccgctgatttaatgtcgttaaatgctgcccatgttcc | Amino acid substitution of amino acids 35 and 36 of TasA sequence by alanines using site-directed mutagenesis |
| KD_AA_35-36_antisense | ggaacatgggcagcatttaacgacattaaatcagcggctgctacttttgcatcag | Amino acid substitution of amino acids 35 and 36 of TasA sequence by alanines using site-directed mutagenesis |
| F_A_72 | gtgatccgttattttcagcttggaaatcctttgtcaacttatctcccggc | Amino acid substitution of amino acid 72 of TasA sequence by alanine using site-directed mutagenesis |

| | | |
|---|---|---|
| F_A_72_antisense | gccgggagataagttgacaaaggatttccaagctgaaaataacggatcac | Amino acid substitution of amino acid 72 of TasA sequence by alanine using site-directed mutagenesis |
| E_A_82 | attaagcgccattagaactgctttgatcgcaagtgatcc | Amino acid substitution of amino acid 82 of TasA sequence by alanine using site-directed mutagenesis |
| E_A_82_antisense | ggatcacttgcgatcaaagcagttctaatggcgcttaat | Amino acid substitution of amino acid 82 of TasA sequence by alanine using site-directed mutagenesis |
| G_A_96 | ggagatgtattgctgccggcgtttgctttaaaatctc | Amino acid substitution of amino acid 96 of TasA sequence by alanine using site-directed mutagenesis |
| G_A_96_antisense | gagattttaaagcaaacgccggcagcaatacatctcc | Amino acid substitution of amino acid 96 of TasA sequence by alanine using site-directed mutagenesis |
| D_A_64 | ggaaatcctttgtcaacttagctcccggctttagatttgat | Amino acid substitution of amino acid 64 of TasA sequence by alanine using site-directed mutagenesis |
| D_A_64_antisense | atcaaatctaaagccgggagctaagttgacaaaggatttcc | Amino acid substitution of amino acid 64 of TasA sequence by alanine using site-directed mutagenesis |
| op-tasA_fw | gctcggtacccggggatcctGCTATAAGGATCAAATGAAATCGTC | Fusion of TasA to mCherry to generate strains JC209, JC222 and JC224 |
| op-tasA_rv | cgccggccatATTTTTATCCTCGCTATGCG | Fusion of TasA to mCherry to generate strains JC209, JC222 and JC224 |
| mCherry_fw | ggataaaaatATGGCCGGCGTTAGCAAAG | Fusion of TasA to mCherry to generate strains JC209, JC222 and JC224 |
| mCherry_rv | actatctaatTTAACCGGTTTTATACAGTTCATCC | Fusion of TasA to mCherry to generate strains JC209, JC222 and JC224 |

| | | |
|---|---|---|
| tasA_downstream_fw | aaccggttaaATTAGATAGTGAATGGGAGAAATTG | Fusion of TasA to mCherry to generate strains JC209, JC222 and JC224 |
| tasA_downstream_rv | tgcatgcctgcaggtcgactATAACAGCAAAAAAAAGAGACG | Fusion of TasA to mCherry to generate strains JC209, JC222 and JC224 |
